# Multi cell line analysis of lysosomal proteomes reveals unique features and novel lysosomal proteins

**DOI:** 10.1101/2020.12.21.423747

**Authors:** Fatema Akter, Srigayatri Ponnaiyan, Bianca Kögler-Mohrbacher, Florian Bleibaum, Markus Damme, Bernhard Y. Renard, Dominic Winter

## Abstract

Lysosomes play a key role in the regulation of cellular metabolism and are increasingly recognized as highly active and diverse organelles which are involved in a large variety of processes. Their essential role is exemplified by the detrimental consequences of lysosomal malfunction, which can result in lysosomal storage disorders, neurodegenerative diseases, and cancer. Using lysosome enrichment and mass spectrometry, we investigated the lysosomal proteomes of six common cell lines. We provide first evidence for cell-type specific differences of lysosomes on a large scale, showing highly variable levels of distinct lysosomal proteins within one cell type, while others are highly conserved among cell lines. Using stable isotope labelling and bimodal distribution analysis, we identify high confidence lysosomal proteins for each cell line. Multi cell line correlation of these data reveals potential novel lysosomal proteins, and we confirm lysosomal localization for five candidates. All data are available via ProteomeXchange with identifier PXD020600.

## Introduction

Lysosomes, the central lytic organelles of the cell, are responsible for the degradation of a large variety of cellular compounds and the recycling of their building blocks, fulfilling a pivotal function for cellular homeostasis. Organelles and macromolecules, which are delivered to lysosomes through endosomes, phagosomes, and different forms of autophagy, are hydrolysed by ~60 enzymes residing in the lysosomal lumen (***Ballabio, 2016***). Defects in degradation, export, or trafficking of these enzymes frequently result in lysosomal dysfunction, causing so-called lysosomal storage disorders (LSDs), a group of rare inherited diseases with detrimental consequences for the affected patients (***Ballabio & Gieselmann, 2009***). Furthermore, in the recent years, lysosomes have been shown to also play a role in more common diseases like neurodegenerative disorders (***Fraldi et al., 2016***) or cancer (***Davidson & Vander Heiden, 2017***).

In addition to lysosomal hydrolases, >100 lysosomal integral and membrane-associated proteins have been confirmed to date. They are involved in a large range of processes including the transport of molecules across the lysosomal membrane, the fusion with vesicles, nutrient sensing, lysosomal positioning, and interaction of lysosomes with other organelles, e.g. for the exchange of metabolites (***Ballabio & Bonifacino, 2019; Xu & Ren, 2015***). Aside from the mammalian target of rapamycin complex 1 (mTORC1), whose members and functions have been investigated in a variety of studies (reviewed in (***Kim & Guan, 2019; R. E. Lawrence & Zoncu, 2019***)), the function of many lysosomal membrane proteins, and the composition of lysosome-associated complexes, remain insufficiently characterized. It is by now well established, however, that in addition to the degradation of cellular macromolecules, lysosomes are involved in a plethora of other fundamental cellular processes. These include, for example, signalling, energy metabolism, protein secretion, antigen presentation, and plasma membrane repair (***Lim & Zoncu, 2016***), indicating that the number of proteins located in or at the lysosomal membrane may be well beyond those which are so-far experimentally validated. In addition, it is known for a long time that lysosomes can differ between individual cell types, and more recently, it is emerging that lysosomes can be sub-classified into individual populations, differing e.g. in their pH value or their mobility (***Ba et al., 2018; Johnson et al., 2016***).

In order to be able to assess such differences on a global scale, and to identify novel lysosomal proteins, unbiased large-scale approaches play a pivotal role. The method that is currently used most frequently is mass spectrometry based proteomics, as it allows to simultaneously identify and quantify large numbers of proteins in a given sample. A variety of studies have been performed, which investigated lysosomes and lysosomal proteins in HEK293 (***Singh et al., 2020; Thelen et al., 2017; Wyant et al., 2018***), HeLa (***Cardoso et al., 2009; Milkereit et al., 2015***), and MEF (***Ponnaiyan et al., 2020***) cells as well as lysosomes derived from tissue (***Bagshaw et al., 2005; Chapel et al., 2013; Markmann et al., 2017***), and primary cells (***Di Lorenzo et al., 2018; Schmidtke et al., 2019***). While the majority of these studies investigated a biological question, such as the alteration of the lysosomal proteome under pathological conditions (***Di Lorenzo et al., 2018; Schmidtke et al., 2019***), some also focused on the identification of novel lysosomal proteins or the improvement/ comparison of techniques for the enrichment of lysosomes, which is a crucial step for the proteomic analysis of lysosomal proteins due to their low abundance (***Abu-Remaileh et al., 2017; Chapel et al., 2013; Ponnaiyan et al., 2020; Singh et al., 2020; Wyant et al., 2018***).

A common feature of datasets generated from lysosome-enriched fractions with sensitive state of the art mass spectrometry approaches, is that a high number of identified proteins are apparently not lysosomal. In a recent study from our group, for example, >7,000 proteins were identified in samples enriched for lysosomes using superparamagnetic iron oxide nanoparticles (SPIONs) (***Ponnaiyan et al., 2020***), which is >60% of the proteins currently estimated to be expressed on average in in-vitro cultivated cells (***Meier et al., 2018***), making it highly unlikely that all of them are related in one way or the other to lysosomes. Also for other approaches, as e.g. the enrichment of lysosomes by sucrose density gradient centrifugation or immunoprecipitation, high numbers of contaminating proteins (based on their gene ontology (GO) classification) are identified (***Singh et al., 2020***), which are most likely binding unspecifically to the beads or columns utilized for the enrichment of lysosomes.

While it is common practice to assess the likelihood of localization to a specific organelle in differential centrifugation approaches by comparison to other fractions (***Chapel et al., 2013; Della Valle et al., 2011***), to our knowledge, with the exception of one study, which assessed the unspecific binding of non-lysosomal proteins to anti-HA beads by label free quantification (***Wyant et al., 2018***), no attempts have been made to discriminate between lysosome-specific and unspecific binding to the affinity columns/ beads. This leaves the question largely unanswered which of the proteins detected in such experiments are truly lysosomal. Furthermore, the vast majority of studies dealing with the proteomic analysis of lysosomes which were enriched from cell lines, investigated only a single type of cells. Due to the high variability achieved among individual lysosome isolation methods and strategies for sample analysis by mass spectrometry based proteomics, even for the same cell line (***Singh et al., 2020***), a comparison of lysosomal proteomes between different cell lines based on published data is therefore not straightforward.

In the current study, we investigated the lysosomal proteome of six widely used cell lines using lysosome enrichment by SPIONs and mass spectrometry based proteomics (***Thelen et al., 2017***). For three of them, we present the first draft of their lysosomal proteome. By comparison of expression levels for lysosomal proteins of the individual cell lines, we identify cell type-specific expression patterns, revealing lysosomal heterogeneity on a global scale. For the identification of truly lysosomal proteins, we developed a novel approach for the definition of background proteins, based on the combination of SPIONs, SILAC (stable isotope labelling by amino acids in cell culture), and a bimodal distribution model for data analysis. Finally, we show that the reproducibility of protein identification across cell lines correlates with the likelihood of lysosomal localization, proposing potential novel lysosomal proteins, and confirming lysosomal localization for selected candidates.

## Results

### Lysosomal stability and recovery vary between cell lines

For lysosome isolation by SPIONs, it is crucial that the nanoparticles added to the cell culture medium are delivered to lysosomes through the endocytic route. In our hands, individual cell lines required distinct conditions to allow for optimal results. Therefore, we initially established the lysosome isolation parameters for HEK293, HeLa, HuH-7, SH-SY5Y, MEF, and NIH3T3 cells. We chose the SPIONs approach over others, as it allows for the most efficient enrichment of large amounts of lysosomes without loss of lysosome-associated complexes (***Singh et al., 2020***). For each cell line, we adapted conditions for coating of plates, density/ number of cells, foetal calf serum (FCS) content, growth time, and time point of SPIONs addition in order to achieve optimal results (for details see ***Supplementary file 1***).

The enrichment of lysosomes from mammalian cells typically results in high numbers of contaminating non-lysosomal proteins (***Ponnaiyan et al., 2020***). In order to be able to discriminate between lysosome-specific proteins and such binding to the column in an unspecific way, we included differentially SILAC labelled control cells - which do not receive SPIONs - acting as an internal standard. The two populations of cells were combined in equal numbers before lysis. In this setting, the likelihood of retention on the magnetic column for unspecifically binding proteins is irrespective of the presence of SPIONs containing lysosomes. Therefore, for control (C) and SPIONs (S) treated samples, these proteins should be detected with a ratio of S/C=1. Proteins, however, which are localized at the lysosome, and therefore enriched in a specific way, should be detected with a ratio of S/C>1 (***Figure 1A***).

**Figure 1.**
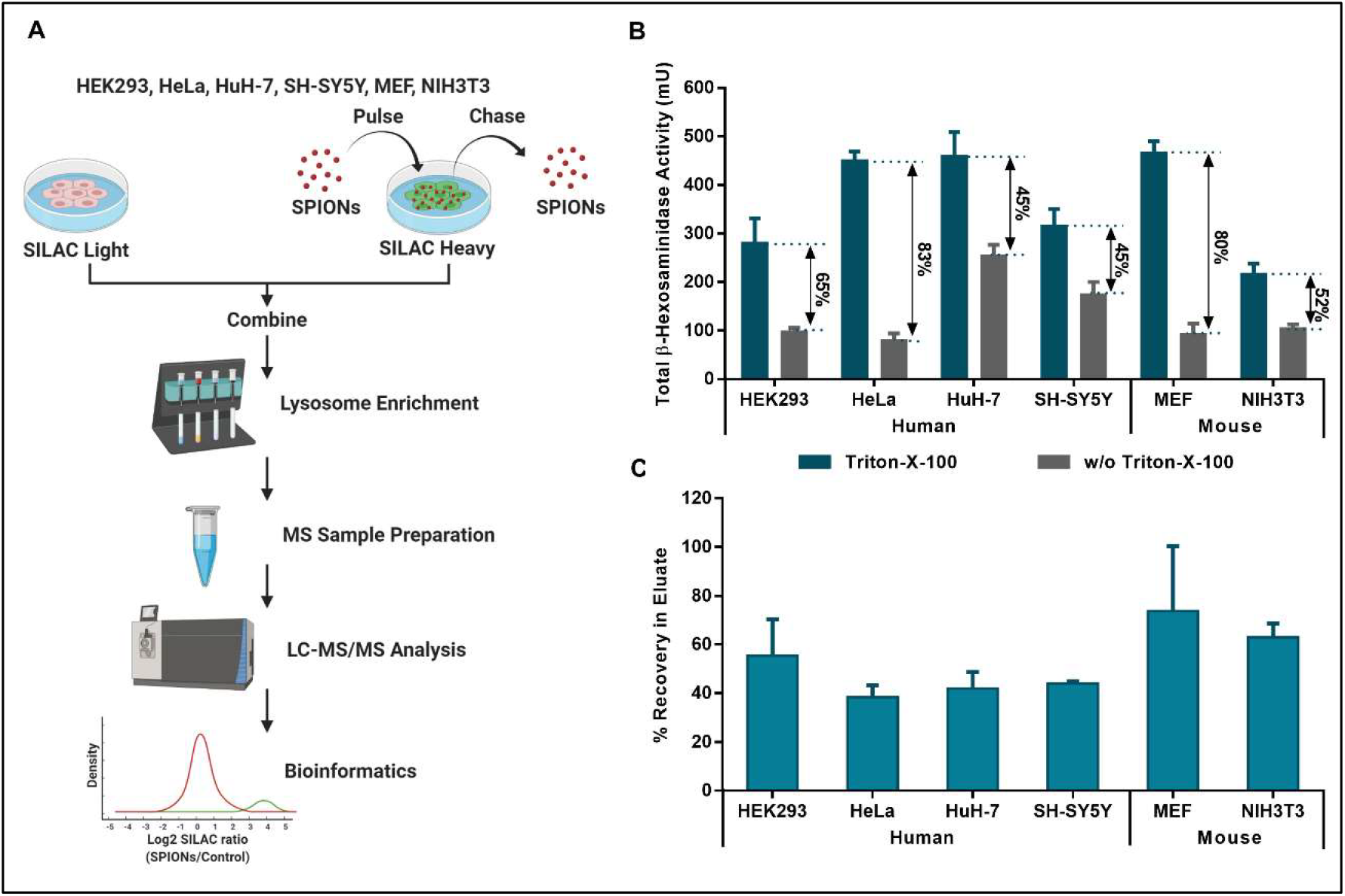
Lysosomal stability and recovery vary across different cell types. **(A)** Workflow for lysosome isolation and mass spectrometric analysis. **(B)** Activity of the lysosomal luminal enzyme β-hexosaminidase determined from post nuclear supernatant fractions of combined light/heavy SILAC cells. β-hexosaminidase activity with/ without the addition of Triton-X-100 relates to the total fraction of lysosomes contained in the sample and those which ruptured during cell lysis, respectively. **(C)** Lysosome recovery rates in input and eluate fractions of lysosome isolation experiments determined by β-hexosaminidase activity. Values are corrected for the presence of differentially SILAC labelled background cells which did not receive SPIONs. SILAC: stable isotope labelling by amino acids in cell culture; SPIONs: superparamagnetic iron oxide nanoparticles; shown are mean values ± SD, n = 4. (***Supplementary file 1***) The following figure supplement is available for figure 1: **Figure supplement 1.** Lysosomal recovery and stability for different cell types.

Following this strategy, we performed four biological replicates for each cell line. In order to account for systematic errors related to SILAC labelling and data analysis, we performed label switching including SPIONs in two replicates of light and heavy labelled cells, respectively. After pooling and disruption of cells, we assessed lysosomal integrity using activity assays for the lysosomal luminal enzyme β-hexosaminidase (***Figure 1B, Figure 1 – figure supplement 1*** (***Thelen et al., 2017***)). The highest total enzymatic activity was determined for MEF, HeLa, and HuH-7 cells, indicating that these cells contain most lysosomes, based on the assumption that the amount of β-hexosaminidase per lysosome is similar across all cell lines (see also ***Figure 4C***). We observed highest percentages of intact lysosomes for MEF and HeLa cells (fraction of intact lysosomes of >80%), while HuH-7 and SH-SY5Y cells revealed highest lability/ organelle disruption (~45% intact lysosomes). For the normalized recovery of intact lysosomes, MEFs performed best, while, surprisingly, HeLa cells yielded the lowest amount, indicating that the percentage of lysosomes receiving SPIONs through unspecific fluid phase endocytosis is lower in these cells compared to e.g. MEFs (~2-fold difference, ***Figure 1 – figure supplement 1, Figure 1C***).

### Proteomic analysis of lysosome-enriched fractions

Using an optimized protocol (***Ponnaiyan et al., 2020***), we analysed the lysosome-enriched fractions of the individual cell lines by mass spectrometry based proteomics (***Figure 1A, Supplementary file 2***). After filtering for peptide and protein identifications with a false discovery rate of 1%, we identified 8,237 proteins from > 1,000,000 peptide spectral matches (PSMs). While the four human cell lines contributed 7,289 unique identifications in total (63%-73% detected in each cell line), 5,235 were identified in the two mouse cell lines with 75% and 90% unique identifications, respectively (***Figure 2A, Figure 2 – figure supplement 1A, Supplementary file 2***). We observed the highest and lowest average numbers of total proteins identified per replicate for HeLa and NIH3T3 cells, respectively, while, when considering only proteins which were covered reproducibly in all four replicates, HeLa, HEK293, and MEF cells performed best (***Figure 2 – figure supplement 1C***).

**Figure 2.**
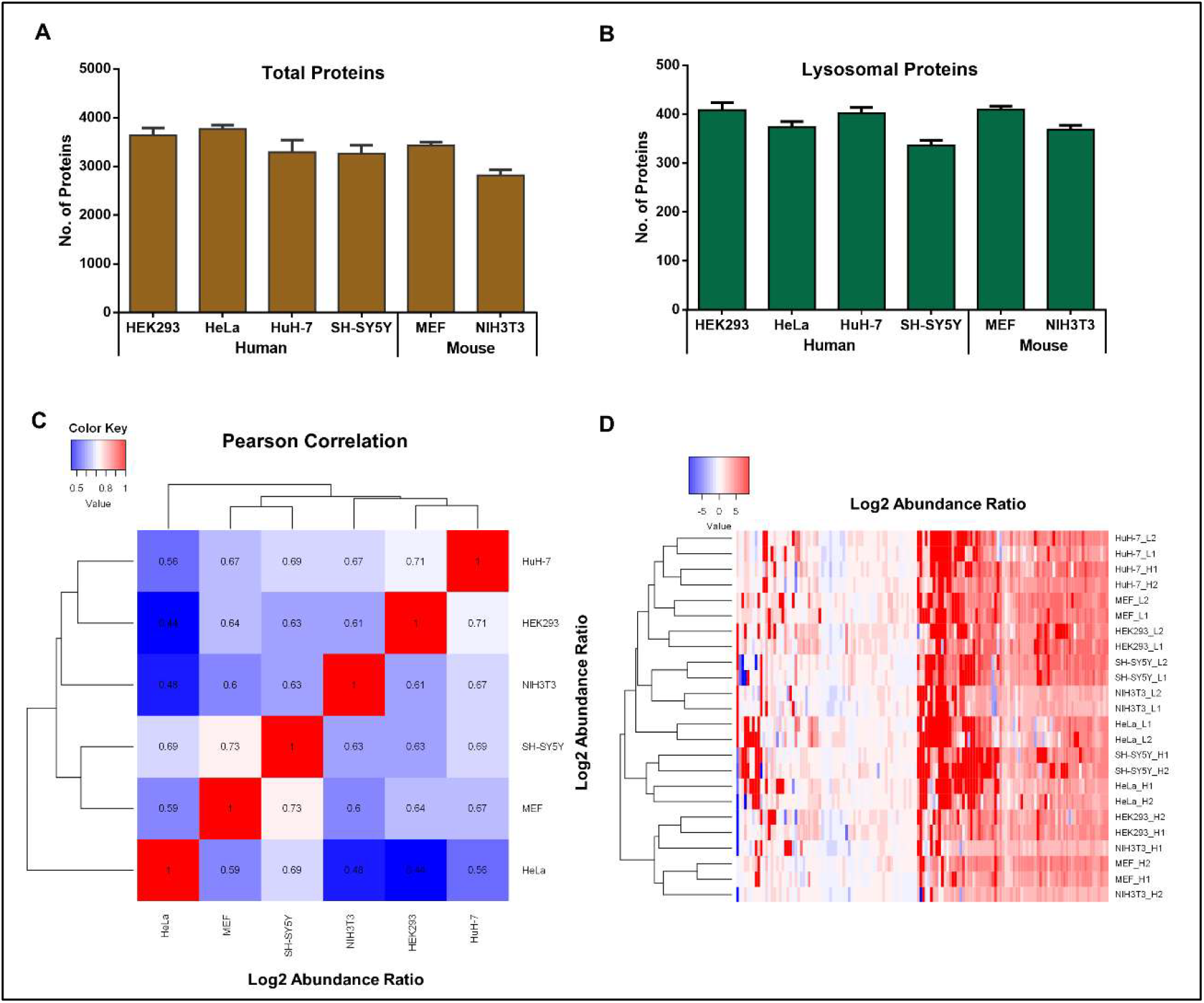
Proteomic analysis of lysosome-enriched fractions. **(A)** Average number of identified proteins detected for individual cell types. **(B)** Average number of identified known lysosomal proteins detected for individual cell types. **(C)** Pearson correlation values of log2 abundance ratios (SPIONs/control) for known lysosomal proteins across individual cell lines. **(D)** Heat map and unsupervised hierarchical clustering of log2 abundance ratios (SPIONs/control) for known lysosomal proteins. SILAC: stable isotope labelling by amino acids in cell culture; SPIONs: superparamagnetic iron oxide nanoparticles; shown are mean values ± SD, n = 4. (***Supplementary file 2***) The following figure supplements are available for figure 2: **Figure supplement 1.** Analysis of lysosome-enriched fractions by mass spectrometry based proteomics. **Figure supplement 2.** Analysis of cell type-specific protein expression profiles based on the datasets from lysosome-enriched fractions of the individual cell lines.

As we were especially interested in proteins located at the lysosome, we subsequently extracted a subset of proteins reported to be located in/ at the lysosome (***Supplementary file 2***). For these known lysosomal proteins, samples derived from HEK293, HuH-7, and MEF cells resulted in the highest average number of identifications for individual runs (***Figure 2B***), with HEK293 cells resulting in superior reproducibility (***Figure 2 – figure supplement 1D***). While SH-SY5Y cells yielded the lowest average number of lysosomal proteins identified per replicate and the lowest contribution to the overall dataset (***Figure 2B, Figure 2 – figure supplement 1B***), they still outperformed both mouse cell lines in terms of reproducibility (***Figure 2 – figure supplement 1D***). In general, even though the numbers of lysosomal proteins identified from all combined replicates was fairly similar between all cell lines, data derived from the two mouse cell lines consistently identified less lysosomal proteins than those from the four human cell lines (***Figure 2 – figure supplement 1D***).

To investigate the correlation among individual cell lines, we determined the Pearson correlation coefficients for the log2 abundance ratios of the identified proteins. On the whole protein level, NIH3T3 cells showed the lowest correlation with all other cell lines, while SH-SY5Y cells were most similar (***Figure 2 – figure supplement 2A***). When considering only known lysosomal proteins, correlation coefficients were in general higher with the exception of HeLa compared to NIH3T3/ HEK293 cells, indicating stronger quantitative differences between the lysosomal proteomes of these cell lines (***Figure 2C***). Based on unsupervised hierarchical clustering, we observed a general clustering for samples which received SPIONs in the light or heavy SILAC channel, respectively (***Figure 2 – figure supplement 2B***). Since especially unspecific binding proteins are affected by the SILAC label switch, this observation underlines the strong contribution of background proteins to the individual datasets. Also for known lysosomal proteins, we observed an effect of the SILAC labelling state of the cell line, but no general clustering occurred dependent on which SILAC label received SPIONs (***Figure 2D***). Intriguingly, irrespective of the analytical approach, and in accordance with the comparison of lysosomal protein identifications (***Figure 2B, Figure 2 – figure supplement 1B, D***), we did not observe any specific trends for human or mouse cells, implying the lack of dominant organism-related properties.

### Correlation of cell type specific protein expression profiles

Based on the heterogeneity of tissue-specific phenotypes for individual lysosomal storage disorders (***Platt et al., 2018***), it is highly likely, that lysosomes in distinct cell types exhibit unique properties. While studies investigating individual lysosomal proteins by e.g. western blot, qPCR, or immunostaining approaches have shown highly variable expression levels across tissues, to our knowledge so far no attempts have been made to compare expression levels of lysosomal proteins between cell lines on a global scale. It remains therefore largely elusive in how far lysosomes differ between cell types.

To further assess differences in the lysosomal proteomes of the individual cell lines analysed, we performed direct comparisons of the identified proteins (***Figure 3, Supplementary file 3***). Out of the 8,704 proteins assigned in total, only 2,173 proteins were found in all six cell lines, while 2,520 were unique to one of them (***Figure 3A***). HEK293 and NIH3T3 cells contributed with 512 and 257 proteins, respectively, the highest and lowest numbers of unique identifications. In all four human cell lines, 326 proteins were identified which were not detected in the two mouse lines, while 235 were unique for the mouse cells.

**Figure 3.**
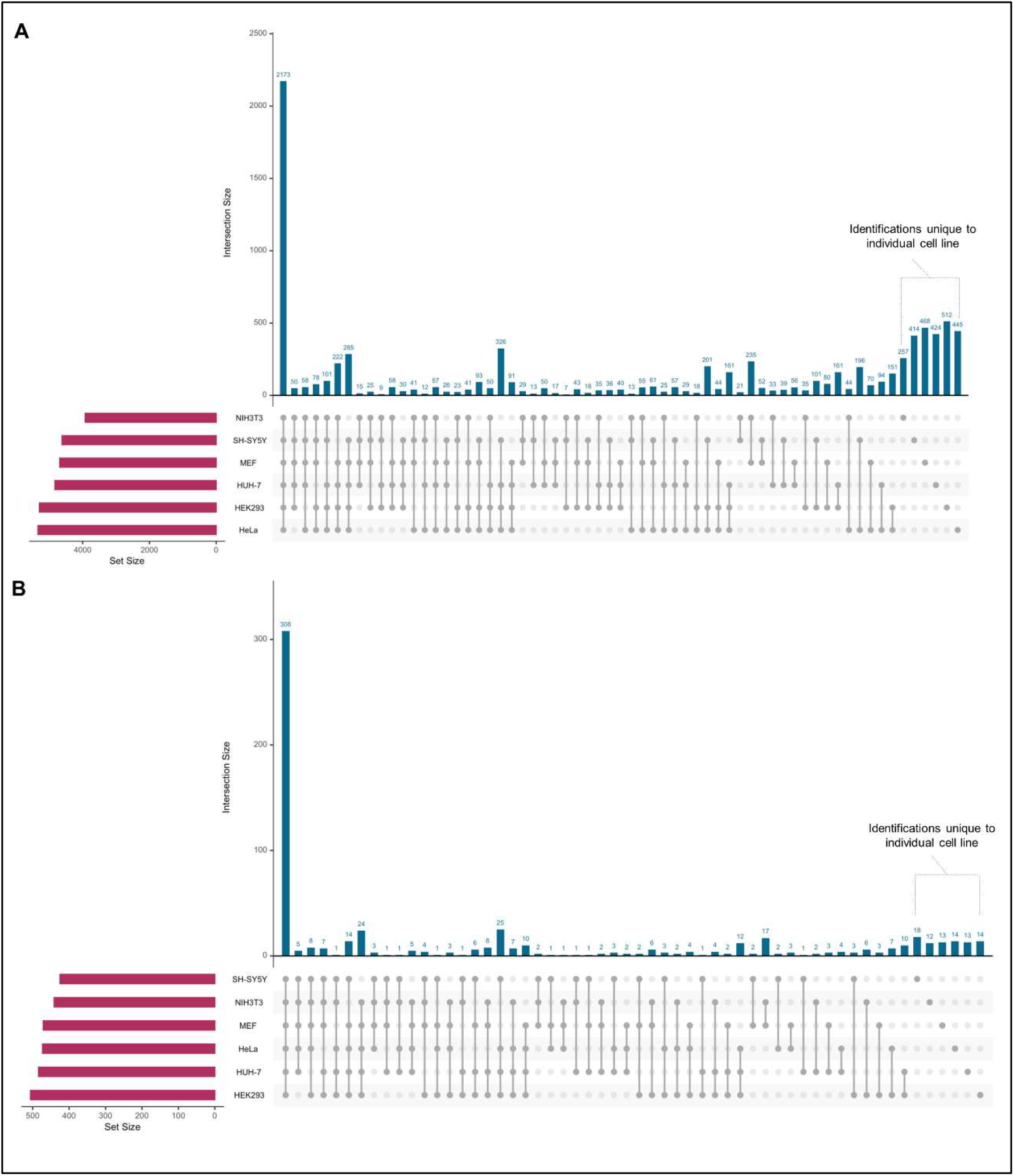
UpSet plots for visualization of overlaps in protein identification for the individual datasets. Dataset size as well as individual overlaps for distinct combinations are indicated (***Supplementary file 3***). **(A)** All proteins identified in the lysosome-enriched fractions of the individual cell types. **(B)** Known lysosomal proteins identified in lysosome-enriched fractions of the individual cell lines.

For the known lysosomal proteins identified across all datasets (643 in total), we observed a similar behaviour, however, with a markedly increased reproducibility compared to the whole dataset (48% of lysosomal proteins vs. 25% of all proteins were identified in all six cell lines, ***Figure 3B***). Furthermore, we detected 130 lysosomal proteins which were unique to human cells (25 reproducibly detected in all four lines), while 42 proteins were unique to mouse cells (17 detected in both cell lines). Overall, while the majority of proteins were reproducibly detected across cell lines, each comparison of the individual proteomes resulted in unique sub-populations. This indicates distinct features of the lysosomes of the respective cells which are possibly related to their individual functions/ characteristics.

### Estimation of abundance levels for known lysosomal proteins

In mass spectrometry experiments, also low abundant proteins are often reproducibly detected, if the sample complexity does not exceed the analytical setup’s limitations defined by the instrument’s sensitivity and speed, as well as the chromatographic gradient length. Therefore, protein identification does not necessarily correlate with abundance, and differences in identification frequently only reveal extreme cases of expression variability, in our case possibly underestimating the extent of lysosomal heterogeneity. Furthermore, every cell requires a certain lysosomal ‘core proteome’ to be able to deal with the turnover of ubiquitous cellular components. We argued that it is therefore highly unlikely that proteins belonging to this class are not expressed at all. It should be more reasonable, that the expression levels of individual proteins belonging to this group are adapted to the specific characteristics of an individual cell type, reflecting its unique composition and function (***Platt et al., 2018***).

To be able to address lysosomal heterogeneity based on changes in protein abundance in more detail, we investigated the expression levels for our list of known lysosomal proteins (***Supplementary file 2***) across all cell lines. We utilized the intensity based absolute quantification (iBAQ) value (***Supplementary file 4***), which is a measure for absolute protein quantity (***Cox & Mann, 2008; Schwanhäusser et al., 2011***), to be able to compare expression levels of human and mouse cells. As the yield of lysosomes varied among cell lines and replicates (***Figure 1 – figure supplement 1***), we corrected for inconsistencies in sample preparation by replicate-wise normalisation of individual iBAQ values (***Supplementary file 5***). For this purpose, we utilized the median iBAQ value from eight core subunits of the V-ATPase complex (detected with ≥ 10 unique peptides each), which is responsible for lysosomal acidification (***Maxson & Grinstein, 2014***), as these proteins were among the most stable among all lysosomal proteins in the datasets (***Figure 4 – figure supplement 1A, Figure 4 – figure supplement 2A***), presenting a ‘core feature’ of lysosomes. This is probably also related to the fact, that the presence and correct stoichiometry of the V-ATPase is essential for lysosomal function. Subsequently, we filtered for proteins with iBAQ values in at least two cell lines and three biological replicates each, calculated the median of the V-ATPase-normalized value for each individual protein, and grouped them into 32 classes based on protein function and/ or localization (***Supplementary file 5***).

### Cell line specific differential expression of lysosomal proteins

Initially, we calculated the sum of all V-ATPase-normalized iBAQ values in the individual cell lines to estimate the overall abundances of lysosomal proteins (***Figure 4 – figure supplement 1B***). Lysosomes enriched from HuH-7 and HeLa cells contained the highest amount of lysosomal proteins (4- and 2.3-fold more than e.g. MEFs, respectively), while values in the other cell types were roughly similar. The observed differences were mainly due to the strong overrepresentation of hydrolases in HuH-7 cells, transporters in HeLa cells, and membrane proteins in both of them. Only lysosome-associated ubiquitin ligases and proteins related to mTORC1, which were most abundant in SH-SY5Y and HEK293 cells, respectively, were highest abundant in other cell lines than HeLa and HuH-7.

Subsequently, we investigated individual lysosomal proteins for their expression levels in the different cell lines. For the V-ATPase complex itself, we detected a conserved stoichiometry for proteins belonging to both the V0 and the V1 part of the complex with the exception of the associated proteins TCIRG1 and ATP6AP1/ ATP6AP2 (***Figure 4 – figure supplement 2A***). For proteins involved in the translocation of small molecules across the lysosomal membrane (***Figure 4A***), we observed a dynamic range of three orders of magnitude with the cholesterol transporter SCARB2 and the Cl^−^/ H^+^ exchanger CLCN6 showing the highest and lowest median expression levels, respectively (difference of ~1,400 fold).

**Figure 4.**
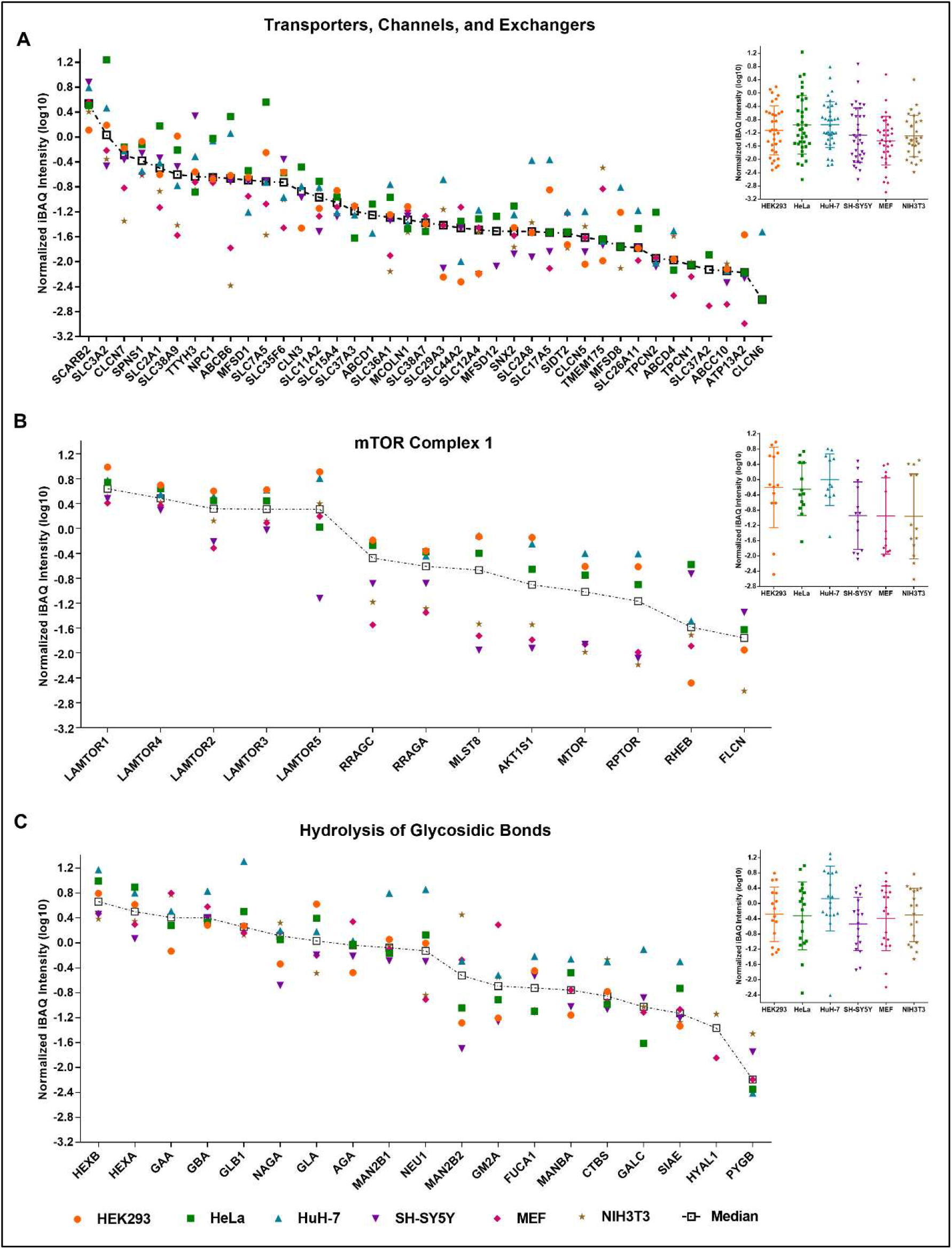
Comparison of expression levels for known lysosomal proteins in individual cell lines. For each protein, the median iBAQ values were determined and normalized to the median intensity of the same eight V-ATPase complex subunits in a replicate-wise manner (***Supplementary file 4, 5***). Proteins are either sorted based on their median intensity (scatter plot) or grouped in a cell line-wise manner (dotted box plot). **(A)** Proteins with known function as transporter, channel, or exchanger. **(B)** Members of, or proteins related to, mTORC1. **(C)** Proteins with a known function related to the hydrolysis of glycosidic bonds. Shown are log10 converted median-normalized iBAQ values for proteins detected in ≥ 3 replicates in each of ≥ 2 cell lines. iBAQ: intensity Based Absolute Quantification. The following figure supplements are available for figure 4: **Figure supplement 1.** Analysis of protein abundance levels by intensity Based Absolute Quantification (iBAQ). **Figure supplement 2.** Individual V-ATPase normalized iBAQ values for proteins grouped in categories of similar function.

Within certain functionally related groups, we detected both highly conserved and highly variable expression patterns. The Ca^2+^ channels MCOLN1, TPCN1, and TPCN2, for example, were expressed at very similar levels in all cell lines, while the amino acid and oligopeptide transporting members of the solute carrier (SLC) family showed both similar and highly variable expression patterns. SLC3A2 and SLC7A5, which are known to heterodimerize (***Alfarsi et al., 2020***), exhibited the highest dynamic range of all proteins in this group with differences of up to 135-fold between the highest (HeLa) and lowest (NIH3T3) expressing cell line, respectively. On the other hand, the sodium dependent amino acid transporter SLC38A7, which was shown to be essential for the extracellular protein dependent growth of cancer cells (***Verdon et al., 2017***), was with a dynamic range of protein expression of ~2-fold most stable. Another member of the SLC38 family, SLC38A9, which is essential for the amino acid dependent activation of mTORC1 (***R. E. Lawrence & Zoncu, 2019***), exhibited with up to ~40-fold a substantially higher difference in expression between cell lines. It was expressed highest in HEK293 cells, correlating with the pattern we observed for other proteins related to mTORC1 (***Figure 4B***). Overall, amino acid/ oligopeptide transporters were expressed especially by HeLa cells at high levels (33-fold higher summed signal intensity compared to NIH3T3 cells), implying a high importance of amino acids derived from lysosomal protein degradation for their metabolism. This is in line with studies reporting the dependence of cancer cell metabolism on amino acids originating from lysosomal degradation of proteins (***Elliott et al., 2019; Verdon et al., 2017***).

For other proteins related to mTORC1, we detected varying patterns (***Figure 4B***). The Ragulator members LAMTOR1-4 were similarly expressed in all cell lines, while LAMTOR5 was ~100-fold more abundant in HEK293 compared to SH-SY5Y cells. For all proteins of this group, we observed, in general, high expression levels in HEK293, HeLa and HuH-7 cells, with the exception of RHEB, which showed for HEK293 cells an inverse behaviour compared to SLC38A9. In accordance with the role of FLCN in the mTORC1 dependent phosphorylation and cytosolic retention of the lysosomal transcription factors TFEB and TFE3 (***Rosalie E Lawrence et al., 2019; Petit et al., 2013***), we either detected FLCN (all cell lines except MEFs) or TFEB/ TFE3 (only MEFs) in the lysosome enriched fractions.

For membrane and membrane-associated proteins, we observed a similar dynamic range as for transporters (***Figure 4 – figure supplement 2B***), with Lamp1/ Lamp2 as highest abundant members. Proteins involved in lysosomal positioning (mainly components of the BORC complex (***J. Pu et al., 2015***)), were present in similar abundances within the respective cell lines, with the exception of ARL8B and DTNBP1 (***Figure 4 – figure supplement 2C***). For lysosome-associated ubiquitin ligases, on the other hand, no trend with respect to certain cell lines was detectable (***Figure 4 – figure supplement 2D***). For hydrolases, the dynamic range of expression was in general smaller than for proteins located at/ in the lysosomal membrane, while proteins related to lipid metabolism (***Figure 4 – figure supplement 2E***) were less variable compared to proteases (***Figure 4 – figure supplement 2F***) and those involved in the hydrolysis of glycosidic bonds (***Figure 4C, Figure 4 – figure supplement 2G***). For the latter, we observed, for example, for GBA highly conserved abundances across cell lines, while other proteins, such as NEU1 or MAN2B2 presented with a variability of up to ~140-fold. For the majority of hydrolases, HuH-7 cells showed highest expression levels, indicating a higher throughput of substrates in their lysosomes, which is probably related to the liver’s function as the main recycling organ of the body.

### Identification of specifically enriched proteins by bimodal distribution analysis

In experiments dealing with the analysis of lysosome-enriched fractions by LC-MSMS, significantly more proteins are identified than those likely to be located at the lysosome, irrespective of the enrichment method or analytical strategy (***Ponnaiyan et al., 2020; Singh et al., 2020; Wyant et al., 2018***). Also in our dataset, we identified for each individual cell line > 4,000 unique proteins in the lysosome-enriched fraction. Based on the currently known group of lysosomal proteins (***Supplementary file 2***), and gene ontology (GO) analyses of the respective datasets (***Mi et al., 2019; Singh et al., 2020; Thomas et al., 2003***), it can be reasonably assumed, that a large portion of these proteins is retained at the column material due to unspecific binding to the stationary phase, or to structures interacting with lysosomes (such as the cytoskeleton), representing contaminations. Therefore, it is difficult to identify novel lysosomal proteins, and studies dealing with the analysis of lysosome-enriched fractions by mass spectrometry based proteomics often do not attempt to infer lysosomal localization from identification, but rather focus on known lysosomal proteins contained in the dataset (***Di Lorenzo et al., 2018; Schmidtke et al., 2019***).

We argued, that inclusion of a population of control cells which did not receive SPIONs should allow for discrimination of lysosome-specific and background proteins, as for unspecific interactors, the chance of enrichment should be irrespective of the presence of SPIONs in the sample. Therefore, in all lysosome enrichment experiments, we combined SPIONs receiving cells with a differentially SILAC labelled population of untreated ‘background cells’ (***Figure 1A***). Based on the signal intensities of the individual SILAC channels, we calculated the log2 transformed ratios of SPIONs/ control, median-normalized the values for individual replicates to compensate for differences in lysosome isolation efficiency, and performed a bimodal distribution analysis. For each cell line, we estimated the mixture of two univariate normal distributions, and assigned a posterior probability of lysosomal localization to each protein using an expectation maximization algorithm (***Dempster et al., 1977***). This resulted in the identification of two overlapping normal distributions presenting on the one hand the ‘background population’ of proteins, which are binding unspecifically to the beads, and on the other hand the ‘lysosomal population’ of proteins, showing increased binding due to the presence of SPIONs in the cells’ lysosomes (***Figure 5, Supplementary file 6***). Based on these distributions, we applied an adjusted p-value cut-off of <0.05 for the definition of enrichment by SPIONs, resulting for each cell line in a high confidence subpopulation of the proteins identified in the individual lysosome-enriched fractions, encompassing from 25% to 41% of the respective datasets (***Supplementary file 7***).

**Figure 5.**
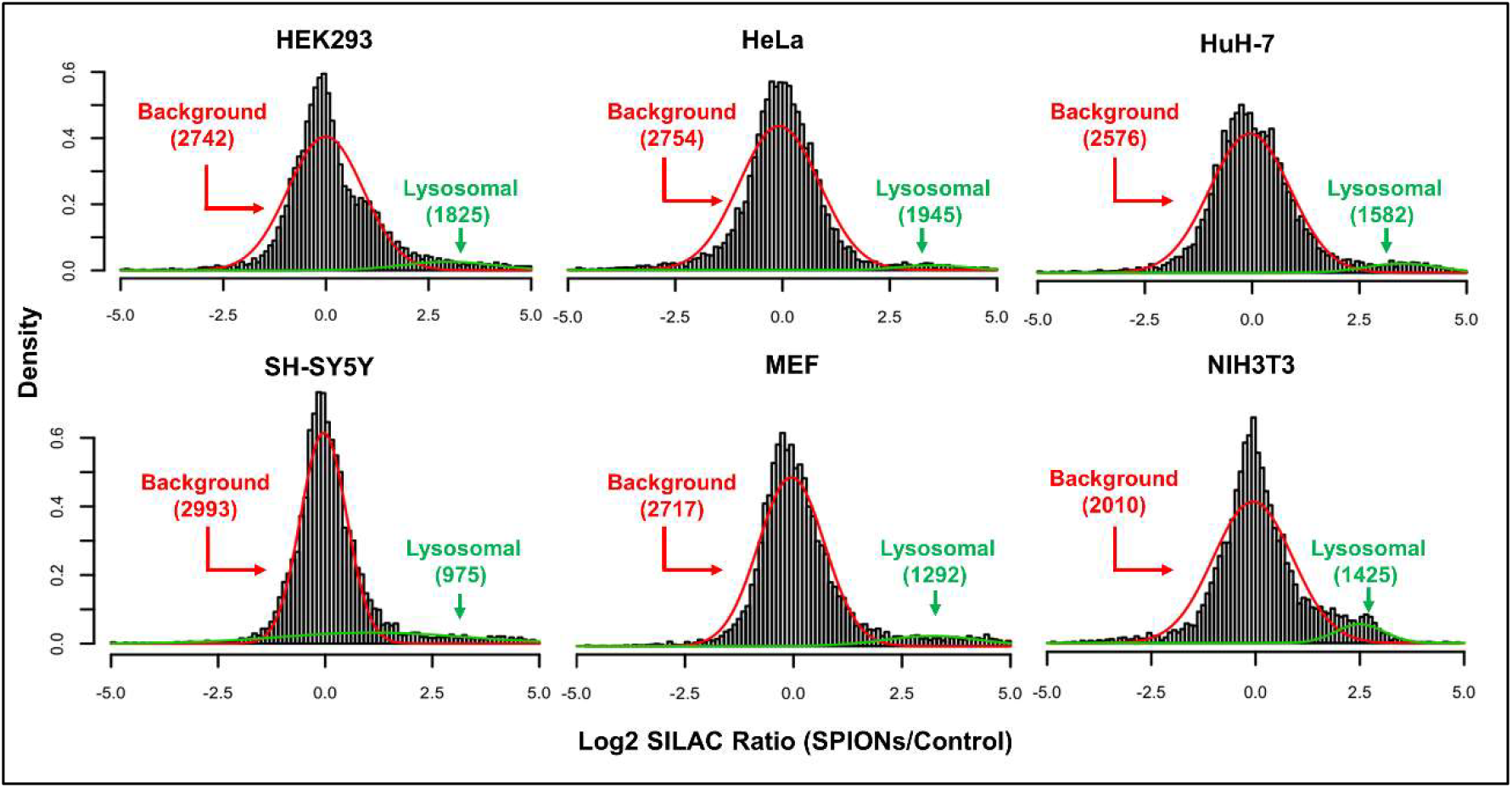
Bimodal distribution analysis of SILAC ratios for proteins quantification values of the individual cell lines. Histograms indicate binned frequencies of log2 transformed normalized SILAC ratios across the datasets. Normal distributed populations were calculated using an expectation-maximization-algorithm and a p-value of ≤ 0.05 was applied as cut-off (***Supplementary file 6, 7***). Red lines indicate the background population showing a similar behaviour between control cells and such receiving SPIONs. Green lines indicate proteins with a significant difference in their SILAC ratio for cells receiving SPIONs relative to the background population. SILAC: stable isotope labelling by amino acids in cell culture; SPIONs: superparamagnetic iron oxide nanoparticles. n = 4.

### Determination of high confidence lysosomal proteins by identification frequency correlation

In order to assess the effect of our bimodal distribution analysis on the individual datasets, we performed GO analyses for all proteins of each dataset, as well as for the sub-populations of high confidence SPIONs-enriched proteins, using the PANTHER overrepresentation test (***Mi et al., 2019; Thomas et al., 2003***). While the proteins in our datasets were assigned to ten cellular compartments before the bimodal distribution analysis, we were able to entirely deplete certain categories - presenting presumably non-specifically enriched proteins - for several cell lines (***Figure 6A, Supplementary file 8***). For example, in HEK293, HuH-7, MEF, and NIH3T3 cells, all nuclear proteins were excluded from the dataset. This was, however, not the case for HeLa and SH-SY5Y cells. We observed a similar effect for proteins related to the cytoskeleton, proteasome, ribosomes, and mitochondria, which were depleted from several cell lines. Most importantly, we were able to achieve an increase for the percentage of known lysosomal proteins in all cell lines, ranging up to 3-fold (in the case of MEFs). This confirms the capability of the approach to enrich the dataset for proteins located at the lysosome. For proteins assigned to endosomes, Golgi apparatus and endoplasmic reticulum, however, only a fraction was removed. Furthermore, compared to the total dataset, the relative abundance of certain contaminating categories further increased in the high confidence SPIONs-enriched proteins, which is due to the reduction in size of the total protein population considered for the analysis.

**Figure 6.**
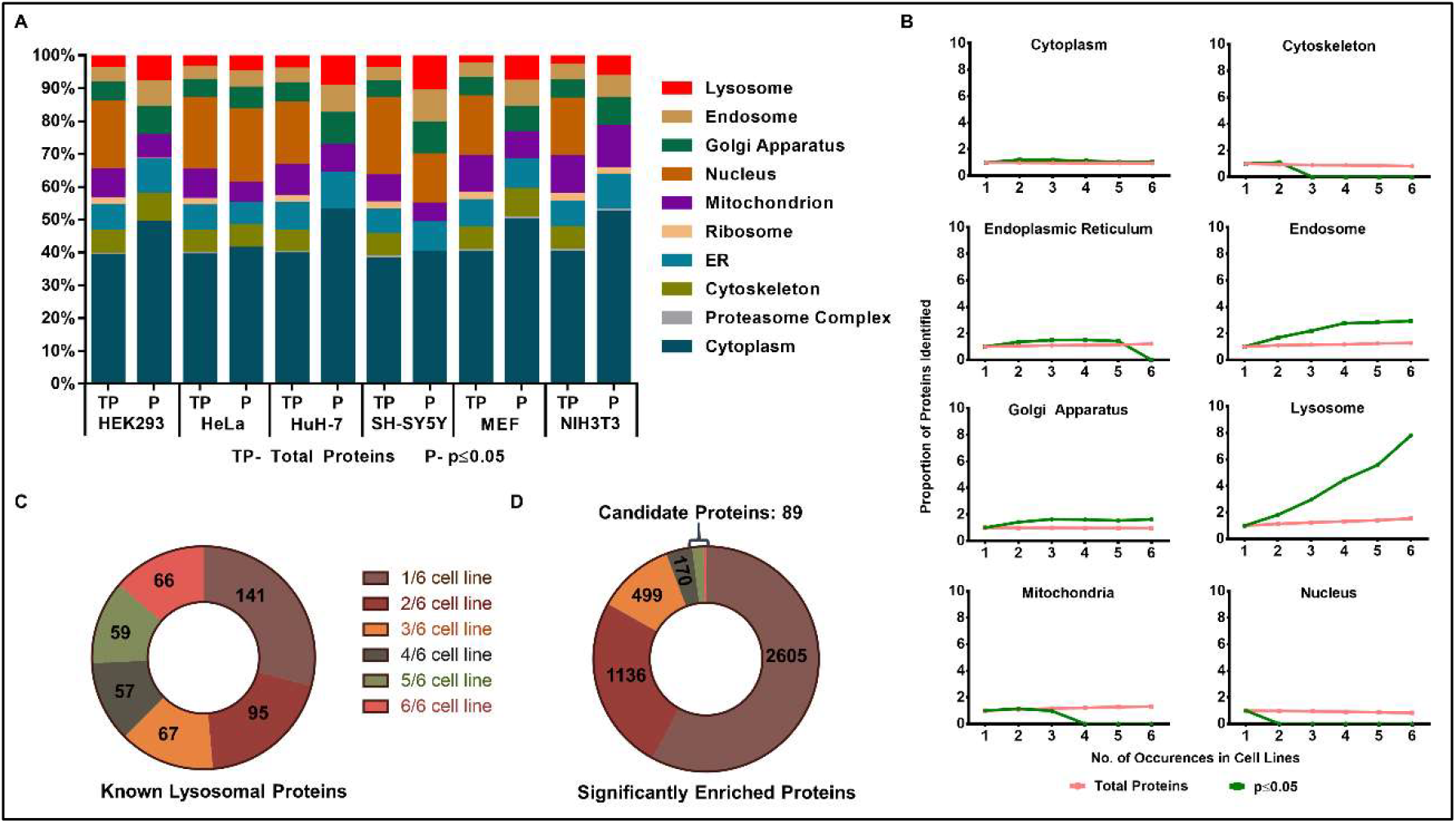
Identification of potential novel lysosomal proteins by analysis of their frequency of detection in multiple cell lines. **(A)** GO analysis of all proteins contained in the respective datasets (total proteins, TP) and proteins which were determined to be significantly overrepresented in SPIONs receiving cells (based on bimodal distribution analysis, p-value ≤ 0.05). The percentage of proteins (relative to the respective dataset) is shown based on their assignment to significantly enriched GO terms (FDR < 0.05, Fisher’s test) (***Supplementary file 8***). **(B)** Correlation of identification frequency and GO term distribution for total proteins and such overrepresented in SPIONS receiving cells. Shown values represent the percentage of proteins assigned to a respective category normalized to the value for considering presence in at least one cell line (***Supplementary file 9***). **(C)** Distribution of known lysosomal proteins dependent on their identification frequency. **(D)** Distribution of proteins determined to be specifically enriched by SPIONs in bimodal distribution analyses excluding known lysosomal proteins dependent on their identification frequency. SPIONs: superparamagnetic iron oxide nanoparticles; GO: Gene Ontology; FDR: False Discovery Rate; n = 4. The following figure supplement is available for figure 6: **Figure supplement 1.** Gene Ontology analysis of high confidence potential novel lysosomal proteins. Also refers to ***Supplementary file 9***

Despite the increase in abundance for known lysosomal proteins, their percentage in relation to the whole dataset peaked at only 11% (SH-SY5Y cells). As we detected known lysosomal proteins more reproducibly across the individual cell lines (***Figure 3B***), we hypothesized that the frequency of identification in multiple cell lines could be used as an indicator for the likelihood of lysosomal localization. We tested this hypothesis by correlation of the percentage of proteins belonging to a certain GO category with their frequency of identification across the individual datasets of the six cell lines (***Figure 6B, Supplementary file 9***). For the whole datasets, we did not observe any correlation of reproducibility with the abundance of a certain category, with the exception of ribosomal proteins, whose percentage increased by 2-fold when comparing proteins identified in 6/ 6 cell lines to those identified in ≥ 1 cell line (***Supplementary file 9***). When we considered our list of p-value filtered high confidence lysosomal proteins, however, we observed a striking increase in the enrichment of lysosomal proteins by up to ~8-fold (proteins identified 6/ 6 vs. ≥ 1 cell lines, ***Figure 6B***). At the same time, we were able to deplete proteins from the cytoskeleton, nucleus, and mitochondria, while the percentage of cytoplasmic and Golgi apparatus proteins remained stable. For proteins assigned to endosomes, we observed the highest increase (~3-fold) after such of lysosomal origin, indicating a specific enrichment, probably due to SPIONs which did not transfer to the lysosomal compartment during the chase period.

The increase in percentage of lysosomal proteins relative to the whole dataset confirmed, that true positive lysosomal proteins are more likely to be reproducibly detected across several cell lines. We further investigated the overlap of known lysosomal proteins with those detected in 5/ 6 or 6/ 6 cell lines and found that 42% and 77% of these populations, respectively, were known lysosomal proteins (***Figure 6C***). Finally, after subtraction of known lysosomal proteins from the list of high confidence lysosomal identifications, we assigned proteins in individual groups based on their identification frequency across cell lines (***Figure 6D***). Out of 4,499 proteins identified to be enriched in at least one of the 6 cell lines, only 2% were reproducibly identified in ≥ 5 cell lines, presenting high confidence potential novel lysosomal proteins (***Table 1***). Among these 89 proteins, GO analysis revealed an enrichment in membrane proteins and transporters (***Figure 6 – figure supplement 1, Supplementary file 9***).

**Table 1.** Potential novel lysosomal proteins based on their overrepresentation in SPIONs receiving cells for ≥ 5 cell lines. Candidates evaluated by immunostaining are highlighted. * TSPAN3 was identified in 6 cell lines, but only enriched in the SPIONs receiving fraction of 4 cell lines.

| Gene Name | # Cell Lines | Gene Name | # Cell Lines | Gene Name | # Cell Lines |
| --- | --- | --- | --- | --- | --- |
| ABHD17B | 6 | DCTN6 | 5 | SLC12A9 | 5 |
| FRRS1 | 6 | DHTKD1 | 5 | SLC2A13 | 5 |
| IFNGR1 | 6 | EIF2B5 | 5 | SLC30A1 | 5 |
| LEPROT | 6 | EPS15 | 5 | SLC33A1 | 5 |
| MFGE8 | 6 | GMPR2 | 5 | SLC38A2 | 5 |
| NDFIP1 | 6 | GOLGA7 | 5 | SLC39A8 | 5 |
| NDFIP2 | 6 | ITFG1 | 5 | SLC46A1 | 5 |
| PIP4P2 | 6 | ITM2B | 5 | SLC4A2 | 5 |
| PLEKHB2 | 6 | LGALS3BP | 5 | SMPD4 | 5 |
| PLPP3 | 6 | LRP10 | 5 | SNX11 | 5 |
| SLC2A6 | 6 | LRP6 | 5 | SNX33 | 5 |
| SLC30A7 | 6 | MFN1 | 5 | STX6 | 5 |
| SLC31A1 | 6 | MMS19 | 5 | TAX1BP1 | 5 |
| TMEM120A | 6 | MPZL1 | 5 | TBC1D15 | 5 |
| TMEM50A | 6 | NIF3L1 | 5 | TGFBR1 | 5 |
| TSPAN6 | 6 | NKIRAS2 | 5 | TM7SF3 | 5 |
| AAK1 | 5 | NT5C2 | 5 | TMEM184C | 5 |
| AGTRAP | 5 | ODR4 | 5 | TMEM268 | 5 |
| ATP9A | 5 | PLXNA1 | 5 | TMEM63B | 5 |
| BNIP1 | 5 | PNPLA8 | 5 | TNC | 5 |
| BRK1 | 5 | PRSS23 | 5 | TPBG | 5 |
| CAB39 | 5 | PSEN2 | 5 | TSN | 5 |
| CARHSP1 | 5 | PSMD14 | 5 | TSPAN14 | 5 |
| CDK5 | 5 | RAB33B | 5 | TXNL1 | 5 |
| CNNM3 | 5 | REPS1 | 5 | UGCG | 5 |
| COPS5 | 5 | RNF149 | 5 | USE1 | 5 |
| CSK | 5 | SCAP | 5 | WDR91 | 5 |
| CTGF | 5 | SERINC1 | 5 | WWP2 | 5 |
| CYR61 | 5 | SERPINE1 | 5 | XXYLT1 | 5 |
| DCTN4 | 5 | SHC1 | 5 | TSPAN3 | 4* |

### Confirmation of lysosomal localization by immunostaining

In order to confirm the validity of our approach, we selected six proteins, which were identified in 6, 5, or 4 cell lines (***Table 1***), and not previously reported to be localized at the lysosome, for follow up studies. Due to the strong enrichment of membrane proteins in our list of high confidence candidates, we focused on proteins which are localized in or at the lysosomal membrane: 1. TM7SF3 (Transmembrane 7 superfamily member 3), which was shown to maintain protein homeostasis through attenuation of ER stress (***Isaac et al., 2017***); 2. SLC12A9 (Solute carrier family 12 member 9), which belongs to the SLC12 family of electroneutral transporters facilitating the symport of Na^+^/ K^+^ with Cl^−^(***Gagnon & Delpire, 2013***); 3. SLC31A1 (High affinity copper uptake protein 1), which belongs to the SLC31 family of copper transporters (***Petris, 2004***); 4. TMEM63B (CSC1-like protein 2), an osmosensitive Ca^2+^ permeable channel (***Marques et al., 2019***); 5. TSPAN3 (Tetraspanin-3), a member of the tetraspanin superfamily which has been involved in trafficking of membrane proteins (***Tokoro et al., 2001***); and 6. NDFIP2 (NEDD4 family-interacting protein 2), which was reported to be involved in controlling the activity of WW-HECT domain E3 ubiquitin ligases, in particular NEDD4 (***Mund & Pelham, 2009***).

For each candidate, we transiently transfected HeLa cells with an expression vector containing an HA- or MYC-tagged version of the respective cDNA and investigated lysosomal localization by co-immunostaining with an antibody directed against the respective tag and the lysosomal marker protein LAMP2 (***Figure 7***). Five of our six candidate proteins (TM7SF3, SLC12A9, SLC31A1, TMEM63B, and TSPAN3) co-localized with LAMP2, thereby confirming their lysosomal localization. For NDFIP2, however, we only observed a partial co-localization with LAMP2 while co-localization with the early endosomal marker Rab5 was much more pronounced. As we also observed enrichment in endosomal proteins in our dataset, we therefore currently believe that NDFIP2 is rather present in endosomes than in lysosomes.

**Figure 7.**
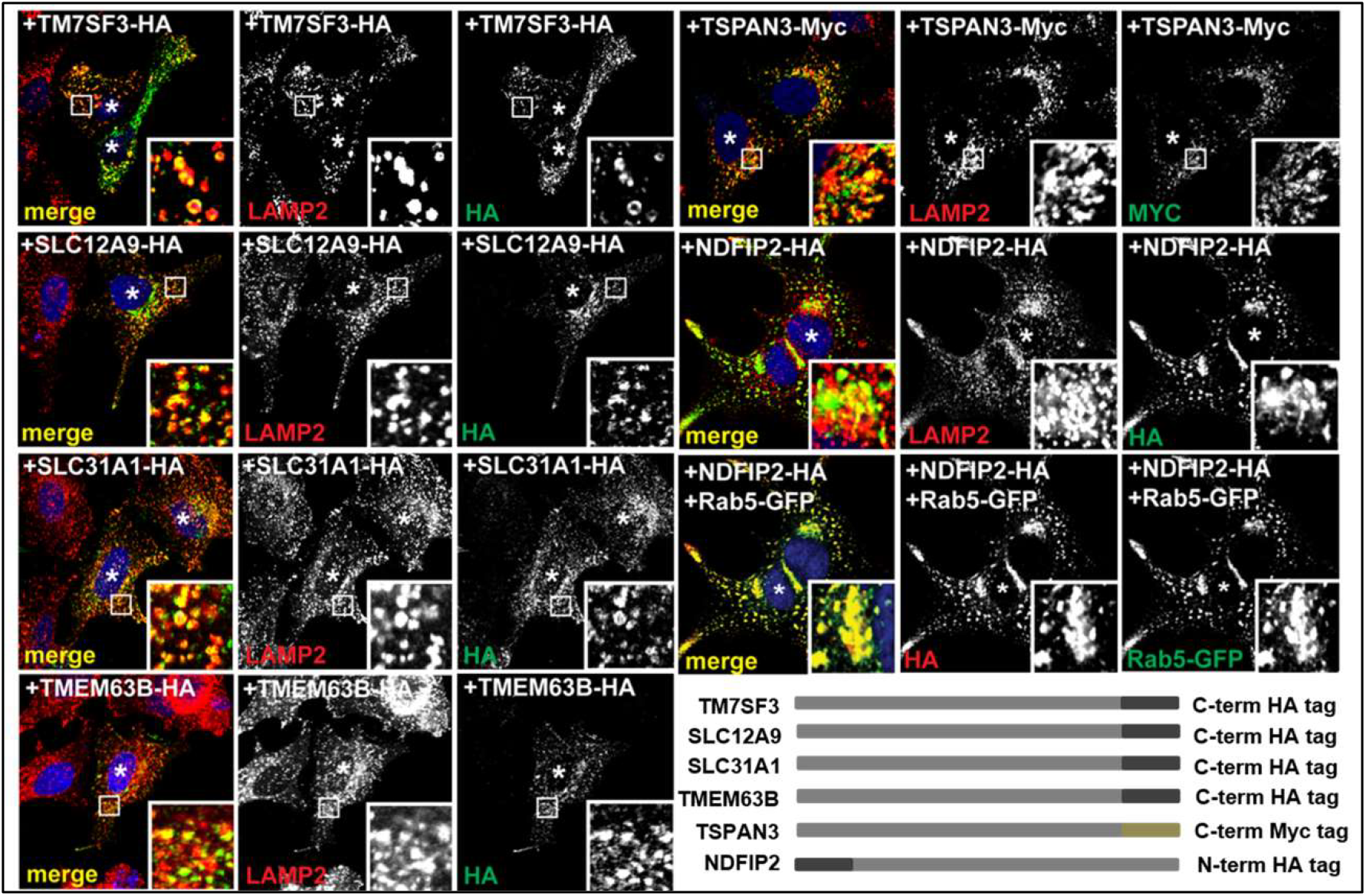
Validation of novel lysosomal proteins by immunostaining in HeLa cells. For six proteins detected in the significantly SPIONs enriched fractions of ≥ 5 cell lines, C-/N-terminally tagged constructs were generated (***Supplementary file 1***). Following transient transfection of HeLa cells, co-localization with lysosomes (Lamp2 staining) or endosomes (Rab5 staining) was investigated by immunostaining.

## Discussion

To our knowledge, this is the first study performing a systematic comparison of lysosome-enriched fractions from different cell lines. Already during cell lysis and lysosome isolation, differences between the individual cell types became apparent, as we observed varying yields of β-hexosaminidase activity and lysosomal intact ratios in the PNS fraction preceding lysosome isolation, as well as different recovery rates in the eluate fraction (***Figure 1B, C***). As the V-ATPase normalized iBAQ levels of β-hexosaminidase were fairly similar across cell lines (***Figure 4C***), the varying yields imply different amounts of lysosomes in these cells. As also the amount of intact lysosomes in the flow through fractions varied considerably (***Figure 1 – figure supplement 1***), these results imply differences in isolation efficiency which may be related to the delivery of SPIONs to the lysosomal compartment. It has to be taken into consideration, however, that we combined SPIONs receiving cells with control cells, which may influence the results.

For the proteomic analysis of lysosome-enriched fractions, we identified for all cell lines fairly similar numbers of proteins. Surprisingly, we did not observe general species-related trends in the global analyses of (lysosomal) proteins (***Figure 2C, 2D, Figure 2 – figure supplement 2A, 2B***), implying no fundamental differences in lysosomal composition between human and mouse cells.

When we investigated protein identification across all cell lines, we detected a highly reproducible core proteome of 2173 proteins, of which 308 have been previously reported to be lysosomal (***Supplementary file 2***). The conserved nature makes these proteins possible key components for lysosomal function. As the comparison of samples based on protein identification does not reflect subtle changes in expression levels, we performed intensity based quantification of our data. To allow for comparison of samples originating from human and mouse cells, which result in different peptides for a given protein, we utilized the iBAQ value, which is an estimate of total abundance (***Schwanhäusser et al., 2011***). We only considered proteins of known lysosomal localization (***Supplementary file 2***) which were reproducibly identified (≥ 3 replicates each for ≥ 2 individual cell lines) and grouped them manually based on their function (***Supplementary file 5***).

As we observed a certain degree of variability between replicates, we normalized each of them on a set of V-ATPase subunits which were detected in all analyses with ≥ 10 peptides. Even though it was shown that the cytosolic V1 part of the V-ATPase is able to dissociate from the V0 part integrated in the lysosomal membrane (***Schwanhäusser et al., 2011***), we did not detect major differences in variability and/ or iBAQs between the subunits of both parts of the complex, implying that it is in its intact state on the SPIONs-enriched lysosomes in our experiments. These data also revealed, that the associated subunits ATP6AP1/ 2 are lower in abundance and more variable across cell lines with the exception of HeLa cells in which they were present at similar abundances (***Figure 4 – figure supplement 2A***).

For the BLOC-one-related complex (BORC), which is involved in microtubule-mediated lysosomal transport (***de Araujo et al., 2020***), we detected most subunits at highly similar levels within a given cell line, but a high variability between cell lines. This implies that the stoichiometry of the complex is conserved, while the number of BORC complexes associating with lysosomes is highly dynamic. A possible reason could be related to differences of lysosomal motility in individual cell types, due to variable demands for lysosomal positioning. Even within single cells, it was shown that lysosomal distribution mechanisms can be heterogeneous, resulting in subpopulations of lysosomes with different moving patterns (***Ba et al., 2018***). Furthermore, we detected a ~8-fold higher abundance of ARL8B relative to ARL8A and the other BORC components. This is probably related to the involvement of ARL8B in other processes, such as e.g. the recruitment of the HOPS tethering complex (***Khatter et al., 2015***), increasing the amount of lysosome-localized ARL8B beyond those associated to BORC.

For mTORC1, we identified distinct patterns across the analysed cell lines. Intriguingly, the members of the Ragulator complex, which consists of LAMTOR1-5 and is responsible for the recruitment of mTORC1 to the lysosomal surface (***R. E. Lawrence & Zoncu, 2019***), were expressed at higher levels than e.g. the RAG GTPases or the mTOR kinase itself. This may be due to Ragulator’s other roles, as e.g. lysosomal positioning, due to its interaction with BORC (***Filipek et al., 2017; Jing Pu et al., 2017***). Along this line, we observed a similar abundance pattern for BORC subunits (***Figure 4 – figure supplement 2C***) and mTORC1 members (***Figure 4B***) in the individual cell lines. For individual members of Ragulator, it was quite surprising to us to find LAMTOR5 at highly variable levels (~100-fold dynamic range), clearly exceeding the dynamic range observed for the other subunits. It was shown that loss of LAMTOR2 induces degradation of the other LAMTOR proteins (***de Araujo et al., 2013***). While these data imply, that correct assembly of Ragulator is crucial for the stability of its members, our data indicate that this may not be the case for LAMTOR5. This would render LAMTOR5 dispensable for the stability of other members of the Ragulator complex, implying a possible function as rate limiting subunit for the full activation of Ragulator (assuming all LAMTORs are needed for Ragulator activity).

For complexes consisting of integral transmembrane transporters and accessory subunits, we found roughly equal stoichiometries confirming their direct relationship to each other. This was, for example, the case for CLCN7 and its β-subunit OSTM1, which are facilitating Cl^−^ import into lysosomes (***Wartosch & Stauber, 2010***). We found for these proteins in all cell lines except NIH3T3, a difference in abundance of < 2-fold. The same applied for ATRAID and SLC37A3, which form a complex releasing nitrogen-containing bisphosphonates from lysosomes (***Yu et al., 2018***). For MFSD1 and its accessory subunit GLMP (***Lopez et al., 2019***), however, we detected a heterogeneous pattern with similar stoichiometry in HeLa and SH-SY5Y cells, while the other cell lines showed a consistently higher abundance of GLMP (up to 22-fold in case of HuH-7 cells), indicating probable other roles for GLMP in certain cell types. Finally, for the complex of the lysosomal cobalamin transporter ABCD4 and the lysosomal membrane protein LMBRD1, which was shown to facilitate the transport of ABCD4 from the ER to lysosomes (***Kawaguchi et al., 2016***), we observed throughout all cell lines a significantly higher abundance of ABCD4 (5 to 26-fold). This indicates that LMBRD1 may be important for the transport of ABCD4, but not for its stability or function, as a roughly similar abundance would have to be maintained in this case.

In general, the iBAQ data should provide a valuable resource to the community, as they, to our knowledge, present the first estimate of the abundance of distinct lysosomal proteins between different cell types. Especially for the conceptualization of experiments, this should allow to choose appropriate model systems, as overexpression of the same protein may result in one cell line in a 2-fold higher abundance, while it will result in a 200-fold increase in another. Furthermore, the iBAQ values allow for an estimate of the importance of certain metabolic pathways in the individual cell lines, as, for example, we observed extremely high levels of most lysosomal enzymes in HuH-7 cells, while concentrations of channels, transporters, and exchangers were rather average.

For the identification of novel lysosomal proteins from our datasets, we applied a novel strategy based on the inclusion of differentially SILAC labelled background cells (which did not receive SPIONs) and bimodal distribution analysis of the resulting samples. While we were able to significantly deplete unspecifically binding proteins from our datasets, and to define a population which was specific for the presence of SPIONs (***Figure 5***), we were not able to fully remove proteins from certain cellular compartments (***Figure 6A***). This is most likely due to the transient interaction of lysosomes with these organelles/ structures, as it is well-established that they form direct contact sites, for example with the endoplasmic reticulum, mitochondria, or the plasma membrane (***Ballabio & Bonifacino, 2019***). As for such cases only a certain number of proteins are probably true interactors which facilitate lysosomal contact, while others are enriched unspecifically through secondary interaction with the respective structure. As such secondary interactions should be less reproducible, we argued that the reproducibility of enrichment, and therefore frequency of identification across the individual datasets, should be reduced for such proteins. Consequently, true lysosomal localization should correlate with reproducibility of identification, which was the case for known lysosomal proteins (***Figure 6B, 6C***). Based on this assumption, we grouped proteins not shown to be localized at lysosomes before and generated a list of potentially novel lysosomal proteins. For reasons of feasibility, we further focused only on such detected in ≥ 5 cell lines for the selection of targets for follow up studies ***Table 1***). Probably also proteins which we identified in ≥ 3 or 4 cell lines present high confidence targets, as we also observed for several known lysosomal proteins a certain heterogeneity of expression (***Figure 3***). Therefore, this list presents in our eyes a valuable resource for future studies investigating novel lysosomal proteins, as it facilitates the assessment of the likelihood of their localization both based on the SILAC ratio relative to control cells and the frequency of detection across the individual datasets.

## Materials and Methods

### Cell culture, stable isotope labelling by amino acids in cell culture (SILAC), and lysosome enrichment

HEK293 (ATCC, CRL-1573), HeLa (ATCC, CCL-2), HuH-7 (JCRB Cell Bank, JCRB0403), SH-SY5Y (ATCC, CRL-2266), MEF (ATCC, CRL-2991), and NIH3T3 (ATCC, CRL-1658) cells were cultured in SILAC DMEM supplemented with 10% dialyzed foetal calf serum (FCS), 100 IU/ml penicillin, 100 μg/ml streptomycin, 181.2 mg/ml light or ^13^C_6_^15^N_2_ lysine (Cambridge Isotope Laboratories, Inc., CNLM-291-H-1), and 87.8 mg/ml light or ^13^C_6_^15^N_4_ arginine (Cambridge Isotope Laboratories, Inc., CNLM-539-H-1) at 37°C, 100% humidity, and 5% CO_2_. Each cell line was passaged at least six times to ensure complete SILAC labelling. For lysosome enrichment of MEF, NIH3T3, HeLa, and HuH-7 cells, 3×10^6^ cells were seeded per 10 cm plate and cultured in DMEM with 2.5% FCS for 72 h. Subsequently, the medium was changed to DMEM with 10% FCS including 10% of SPIONs solution (DexoMAG40, Liquids Research Ltd, BKS25) for either light or heavy SILAC labelled cells (2 replicates each), followed by incubation for 24 h. For HEK293 cells, plates were coated with 100 μg/ml poly-L-lysine (Sigma Aldrich, P1524). For HEK293 and SH-SY5Y cells, 6×10^6^ cells were seeded directly in DMEM with 10% FCS and 10% SPIONs followed by incubation for 24 h. Subsequently, for all cell lines, cells were washed twice with 1x PBS, fresh medium was added, and the cells incubated for 36 h. Cells were washed three times with ice-cold 1x PBS, scraped in 2 mL isolation buffer [250 mM sucrose, 10 mM HEPES/ NaOH pH 7.4, 15 mM KCl, 1 mM CaCl_2_, 1 mM MgCl_2_, 1.5 mM MgAc, 1 mM dithiothreitol (DTT), 1x cOmplete EDTA-free protease inhibitor cocktail (Roche Diagnostics GmbH, 11873580001)] per plate, and lysed using a 15 ml dounce homogenizer. Nuclei and intact cells were pelleted by centrifugation at 4 °C, 500 g for 10 min, and the post-nuclear supernatant (PNS) was transferred to a new tube. The pellet was resuspended in 2 mL isolation buffer, the procedure repeated, and the PNS fractions combined. The pooled PNS fractions of the individual cell lines were passed by gravity flow through LS columns (Miltenyi Biotech, 130-042-401) placed in a MidiMACS Separator (Miltenyi Biotech, 130-042-302), and washed with 5 mL isolation buffer. Lysosomes were eluted from the columns twice with 1 mL isolation buffer each using a plunger. Lysosomal integrity and isolation efficiency were assessed using the β-hexosaminidase assay with/ without the addition of 0.8% Triton X-100 (v/v, final concentration) (***Thelen et al., 2017***). To calculate the relative recovery of lysosomes in the eluate fractions, only the contribution of cell lines receiving SPIONs was considered. The protein concentration was determined using the DC protein assay (BioRad, 5000116).

### Sample preparation for mass spectrometry

For each sample, 100 μg of protein were precipitated by ice-cold chloroform/ methanol (2:1, v/v) as described elsewhere (***Winter & Steen, 2011***). Protein pellets were resuspended in 8 M urea, 0.1 M TEAB (***Kollipara & Zahedi, 2013***), and incubated at room temperature (RT) for 45 min, 800 rpm followed by reduction with 5 mM dithiothreitol (DTT) at 56 °C, 800 rpm for 25 min, and alkylation with 20 mM acrylamide at RT for 30 min (***Muller & Winter, 2017***). The reaction was quenched by addition of 5 mM DTT, samples diluted to 4 M urea with 0.1 M TEAB, rLys-C (Promega, V1671) added at an enzyme to protein ratio of 1 to 100 (w/w), and digestion was performed at 37 °C overnight. The following day, samples were diluted with 0.1 M TEAB to 1.6 M urea, trypsin (Promega, V5111) was added at an enzyme to protein ratio of 1 to 100 (w/w), and the samples were incubated at 37 °C for 10 h. Finally, the samples were acidified using acetic acid (0.1% final concentration) and 10 μg of peptides desalted by STAGE tips as described elsewhere (***Rappsilber et al., 2003***). Eluted peptides were dried using a vacuum centrifuge and resuspended in 5% acetonitrile (ACN)/ 5% formic acid (FA).

### LC-MS/MS analysis

Mass spectrometry analyses were performed using a Dionex Ultimate 3000 system coupled to an Orbitrap Fusion Lumos mass spectrometer (both Thermo Scientific). Columns were manufactured in-house as follows: 50 cm spray tips were generated from 360 μm outer diameter/ 100 μm inner diameter fused silica capillaries using a P-2000 laser puller (Sutter Instruments) and packed with 1.9 μm Reprosil AQ C_18_ particles (Dr. Maisch, Ammerbuch-Entringen, r119.aq). Peptides were resuspended in 5% ACN/ 5% FA and loaded on the analytical column at a flow rate of 600 nL/min, 100% solvent A (0.1% FA in water). Subsequently, the separation was performed at a flow rate of 300 nL/min with a 240 min linear gradient from 5–35% solvent B (95% ACN/ 0.1% FA). Survey spectra were acquired in the Orbitrap mass analyser with a mass range of m/z 375–1,575 at a resolution of 60,000. MS/MS fragmentation was performed in the data dependent acquisition (DDA) mode for charge states between 2–4 by higher-energy collisional dissociation HCD and data were acquired in the Orbitrap at a resolution of 30,000. The cycle time was set to 5 s and the precursor isolation width to 1.6 m/z using the quadrupole. For MS1 and MS2 scans, the automatic gain control (AGC) was set to 4 × 10^5^ and 5 × 10^5^, respectively.

### Mass spectrometry data analysis

Thermo *.raw files were analysed with Proteome Discoverer 2.2 (Thermo Fisher Scientific) in combination with Mascot 2.6.1 (www.matrixscience.com). For database searching, UniProt *Homo sapiens* (release 2019_05, 73,920 entries) and UniProt *Mus musculus* (release 2019_05, 54,425 entries) in combination with the cRAP database (ftp://ftp.thegpm.org/fasta/cRAP/crap.fasta) including common contaminants were used. The following parameters were defined: variable modifications: oxidation of methionine, acetylation of protein N-termini; fixed modification: propionamide at cysteine; mass tolerance: 10 ppm for precursor ions, 50 mmu for fragment ions; enzyme: trypsin except proline was the next amino acid; missed cleavage sites: 2. Data were filtered with a false discovery rate (FDR) of 1% at the peptide level using Percolator (***Spivak et al., 2009***) and proteins were exported with an FDR of 1%. For quantification, SPIONs/ control ratios were determined for each individual raw file and unique as well as razor peptides were used. iBAQ values were calculated using MaxQuant version 1.6.14.0 in combination with the UniProt reference proteomes for *Homo sapiens* (release 2020_04, 96,808 entries) and *Mus musculus* (release 2020_04, 63,666 entries). The same search parameters as for Proteome Discoverer were applied with the exception of definition of heavy labelled arginine (Arg10) and lysine (Lys8) as fixed modification for cells receiving SPIONs in the heavy SILAC channel.

### Bioinformatics analysis

Only peptides identified with high confidence were exported from Proteome Discoverer and MaxQuant for further analysis using R 3.5.1 (2018-07-02) (***Team, 2013***), Microsoft Excel 2016, and GraphPad Prism 6.07 (GraphPad Software). Abundance ratios of SILAC-labelled proteins (SPIONs/ control) were log2-transformed and median-normalized using R. To estimate the parameters of the probability distribution, an Expectation-Maximization-Algorithm (EM algorithm) (***Dempster et al., 1977***) was applied using the normalmixEM function of the R mix tools package (***Benaglia et al., 2009***). Setting the starting values [mu = (0.5) and sigma = (1.1)], estimators for mixtures of two univariate normal distributions were calculated, and a posterior probability was assigned to each protein by inserting the estimated parameters into the formula:

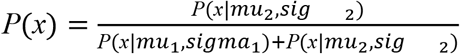

The derived probability describes the likelihood of the observation based on the calculated model. Extreme abundance ratio values (100 and 0.01) were handled separately, as they indicate missing values in one of the distributions and are arbitrarily set by the Proteome Discoverer. Therefore, they were replaced with the maximal or minimal abundance ratios observed during data analysis and the posterior probabilities of these values were set to 1 or 0, respectively. For each dataset, the FDR was calculated applying the Benjamini–Hochberg (***Benjamini & Hochberg, 1995***) multiple testing procedure of R (***Eddelbuettel et al., 2019***) with a default significance level (alpha = 0.05). Principal component analysis (PCA) was performed using the prcomp function of the R package stats, version 3.6.2, setting center = TRUE and scale = TRUE (***Team, 2013***). Pearson correlation coefficients were calculated with the rcorr function from the R package Hmisc, version 4.4-0 (type = “pearson”) (***Harrell Jr, 2019***). Heatmaps of abundance ratios and Pearson correlation were created with the heatmap.2 function from the R package gplots, version 3.0.3, and using hierarchical clustering (order = hclust) (***Warnes et al., 2016***). For UpSet plots, Proteome Discoverer data were joined using the full join function of the R package dplyr, version 0.7.8 (***Hadley Wickham et al., 2017***). If a protein had been found in a cell line, the value was set to 1. Otherwise, if there was a missing value, the respective cell was set to 0. UpSet plots were generated with the upset function from the R package UpSetR, version 1.4.0, setting order.by = degree (***Conway et al., 2017***). For processing of iBAQ values, the median iBAQ intensity of V-ATPase subunits detected with ≥ 10 unique peptides were calculated and used to normalize the other proteins of the respective replicate. Subsequently, the median of the V-ATPase normalized values across individual replicates was calculated and utilized for further analyses. Gene ontology (GO) and functional enrichment analyses were performed using PANTHER (http://pantherdb.org/, database release date 2019-02-02) and g:Profiler (https://biit.cs.ut.ee/gprofiler/) (***Mi et al., 2019; Raudvere et al., 2019***). Over-representation of subcellular localization was assessed with the GO cellular component complete function of PANTHER (p<0.05, Benjamini-Hochberg, FDR corrected). For functional analysis of high confidence potential novel lysosomal proteins, the Gene Group Functional Profiling (g:GOSt) tool available in the g:Profiler web server (version from 2020-03-09) was used. Statistical enrichment analysis with g:GOSt was performed with GO molecular function, GO cellular component, GO biological process and Reactome (p<0.05, g:SCS threshold method).

### RT-PCR and molecular cloning

RNA was isolated from MEFs or NIH3T3 cells using the Nucleospin RNA plus kit (Macherey-Nagel, 740984.5). Subsequently, cDNA was generated using the RevertAid First Strand Synthesis kit (Thermo Fisher Scientific, K1621) according to the manufacturer’s instructions utilizing random hexamer primers. Downstream RT-PCR amplification of lysosomal/ endosomal genes was carried out with the respective set of primers (***Supplementary file 1***). PCR products were purified with the High Pure PCR Product Purification Kit (Sigma Aldrich, 11732676001) and cloned into the pcDNA3.1 Hygro+ mammalian expression vector (Invitrogen, V87020). Successful cloning and sequence identity for the respective gene’s cDNA was confirmed using Sanger sequencing (Eurofins) for each generated plasmid. The Tspan3-myc expression vector was a kind gift from Paul Saftig and the Rab5-GFP expression vector was a kind gift from Sergio Grinstein.

### Immunostaining and confocal laser microscopy

HeLa cells were cultured using cell culture media as described above. Cells were seeded on glass coverslips one day prior to transfection. The transfection was carried out with 750 ng plasmid DNA using the TurboFect transfection reagent (Thermo Fisher Scientific, R0534). After the transfection, all cells were cultured for additional 48 h prior to immunostaining. Transfected HeLa cells were washed 3x with 1x PBS and fixated with either methanol or 4% PFA in 1x PBS for either 10 or 20 min. Subsequently, all samples were rewashed 3x with 1x PBS and incubated for 5 min in 1x PBS with 0.2% saponin and for 15 min in 1x PBS with 0.02% glycine and 0.2% saponin. All samples were blocked for 1 hour in 1x PBS with 0.2% saponin and 10% FCS. After blocking, cells were stained with primary antibodies against HA (3F10, Sigma Aldrich, 12158167001) or Myc (71D10, Cell signaling, 2278) and LAMP2 (H4B4, DSHB, H4B4) overnight at 4 °C. The next day, all samples were repeatedly washed in 1x PBS with 0.2% saponin and incubated with secondary antibodies donkey anti-rat Alexa fluor 488 (Thermo Fisher Scientific, A21208) or donkey anti-rabbit Alexa fluor 488 (Thermo Fisher Scientific, A21206) and donkey anti-mouse Alexa fluor 594 (Thermo Fisher Scientific, A21203) for 1 h at RT. After additional washing in 1x PBS with 0.2% saponin, coverslips were rinsed twice with H2O before embedding them in Mowiol-DABCO with DAPI (4-,6-diamidino-2-phenylindole). Cells were analysed with an inverted confocal laser scanning microscope (FV1000, Olympus). Image acquisition was performed with a UPLSAPO 60x oil immersion objective (NA:1.35) and 2x zoom. All images were acquired and processed with the Olympus FluoView Software.

## Abbreviations

SPIONs: Superparamagnetic Iron Oxide Nanoparticles
SILAC: Stable Isotope Labelling of Amino Acids in Cell Culture
LC-MSMS: Liquid Chromatography-tandem Mass Spectrometry
iBAQ: intensity Based Absolute Quantification

## Data availability

The mass spectrometry proteomics data have been deposited to the ProteomeXchange Consortium via the PRIDE (***Perez-Riverol et al., 2019***) partner repository with the dataset identifier PXD020600.

## Acknowledgments

We thank Volkmar Gieselmann, Robert Hardt, Jasjot Singh, and Paul Saftig for valuable discussions.

## Additional Information

### Funding

This study was supported by the German Academic Exchange Service (DAAD), the National overseas scholarship by the Indian Ministry of Social Justice and Empowerment, and the DFG Research Unit FOR2625.

### Author contributions

FA, SP, and DW designed the experiments. FA and SP performed lysosome enrichment and mass spectrometry-related experiments. FB and MD performed immunostaining experiments. FA, SP, DW, BKM, and BYR analysed the data. DW, FA, and SP wrote the manuscript. All authors edited and approved the manuscript.

### Disclosure statement

The authors declare no conflict of interest.

### Additional Files

Supplementary file 1. Cell culture conditions for lysosome isolation by superparamagnetic iron oxide nanoparticles (SPIONs) from six different cell lines. Primers for cloning of target genes. Refers to ***Figure 1 and Figure 7***

Supplementary file 2. Curated list of known lysosomal proteins (based on Thelen et al. 2017 and the PANTHER/UniProt databases). Proteins identified in proteomic analyses of lysosome enriched samples from six different cell lines. Refers to ***Figure 2, Figure 2 – figure supplement 1, 2***

Supplementary file 3. Reproducibility of protein identification in proteomic analyses of lysosome enriched samples from six different cell lines. Refers to ***Figure 3***

Supplementary file 4. Intensity based absolute quantification (iBAQ) values for the proteomic analyses of lysosome enriched samples from six different cell lines. Refers to ***Figure 4, Figure 4 – figure supplement 1, 2***

Supplementary file 5. V-ATPase normalized iBAQ values for known lysosomal proteins and assignment to individual lysosomal protein classes. Refers to ***Figure 4, Figure 4 – figure supplement 1, 2***

Supplementary file 6. Localization probabilities of individual proteins to the lysosomal fraction determined by bimodal distribution analysis of SILAC intensity ratios. Refers to ***Figure 5***

Supplementary file 7. Putative lysosomal proteins determined by bimodal distribution and multi-cell line correlation analysis. Refers to ***Figure 5***

Supplementary file 8. Cellular component GO analyses using the PANTHER overrepresentation test. Refers to ***Figure 6***

Supplementary file 9. Correlation of distinct GO categories and protein identification frequencies across six cell lines. Gene set enrichment analysis for proteins assigned to the lysosomal compartment by bimodal distribution analysis in ≥ 5 cell lines. Refers to ***Figure 6-figure supplement 1***

## Notes

### Competing Interest Statement

The authors have declared no competing interest.

