## Supplementary figures for "Multi cell line analysis of lysosomal proteomes reveals unique features and novel lysosomal proteins"

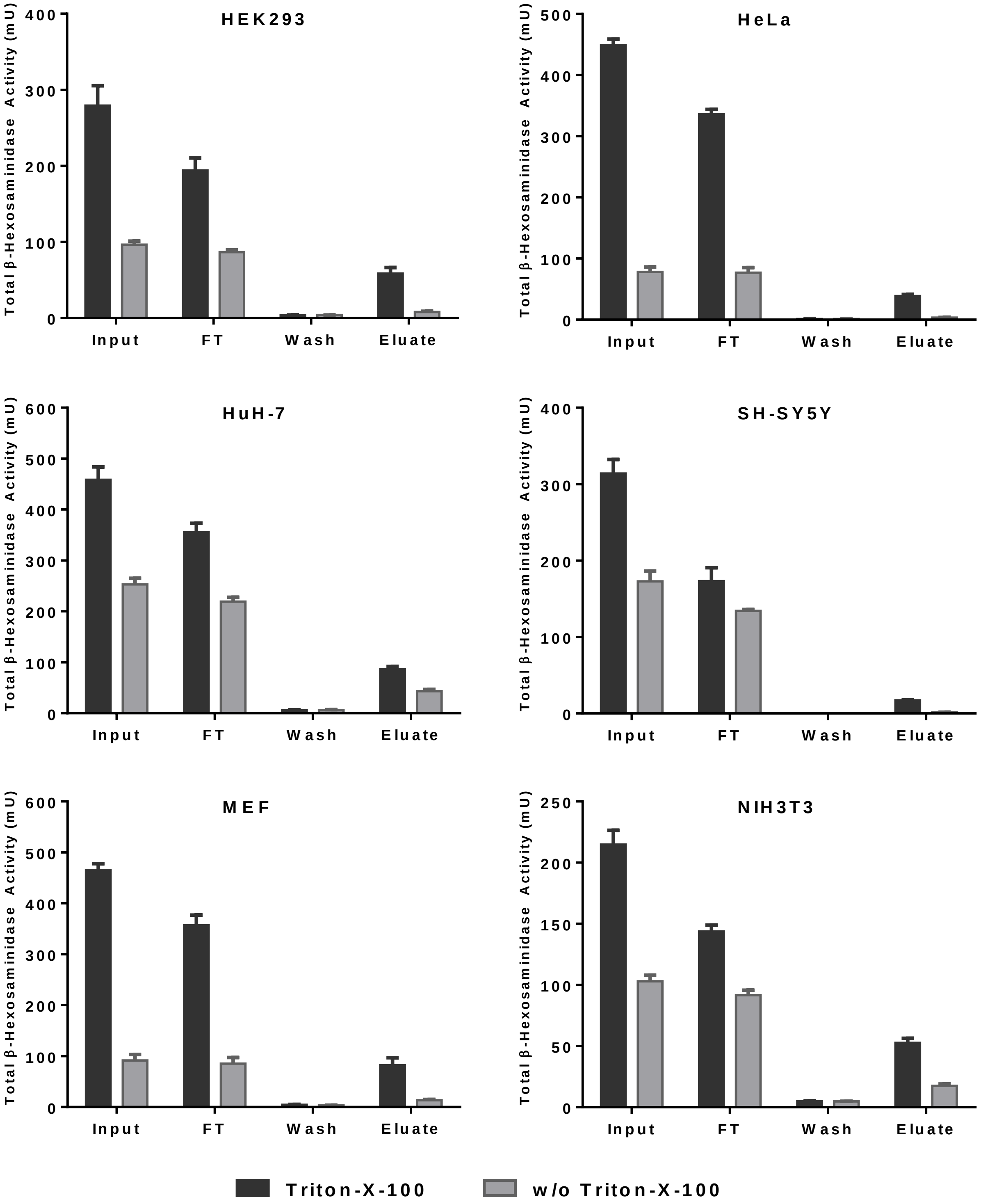


**Figure 1-figure supplement 1: Lysosomal recovery and stability for different cell types.** Shown are total β-hexosaminidase activities determined by enzyme activity assays for individual fractions of SPIONs-based lysosome isolation from a 1:1 mixture of differentially SILAC labelled control and SPIONs receiving cells. Comparison of samples with and without Triton-X-100 allows for estimation of percentage of intact lysosomes. FT: flow through, mU: milli units; SILAC: stable isotope labelling by amino acids in cell culture; SPIONs: superparamagnetic iron oxide nanoparticles. Shown are mean values ± SD, n = 4.

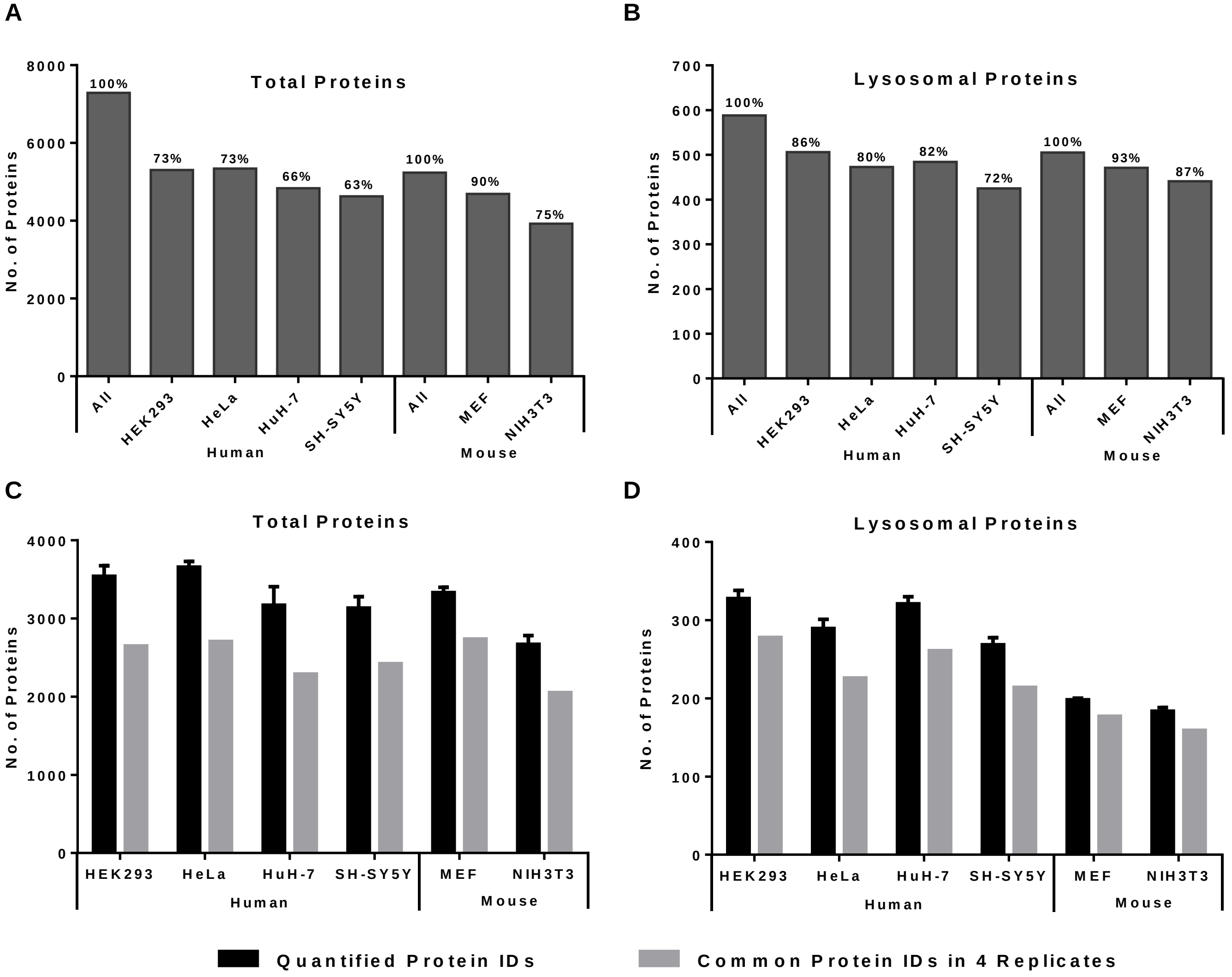


**Figure 2-figure supplement 1: Analysis of lysosome-enriched fractions by mass spectrometry based proteomics.** (**A**) Total protein numbers and identification rates relative to the sum of proteins identified in the whole study for the datasets of the individual cell lines. (**B**) Total numbers and identification rates relative to the sum of known lysosomal proteins identified in the whole study of known lysosomal proteins for the individual cell lines. (**C**) Numbers of quantified proteins and such detected in all four biological replicates for the datasets of the individual cell lines. (**D**) Numbers of quantified proteins and such detected in all four biological replicates for known lysosomal proteins in the individual cell lines. Shown are mean values ± SD, n = 4.

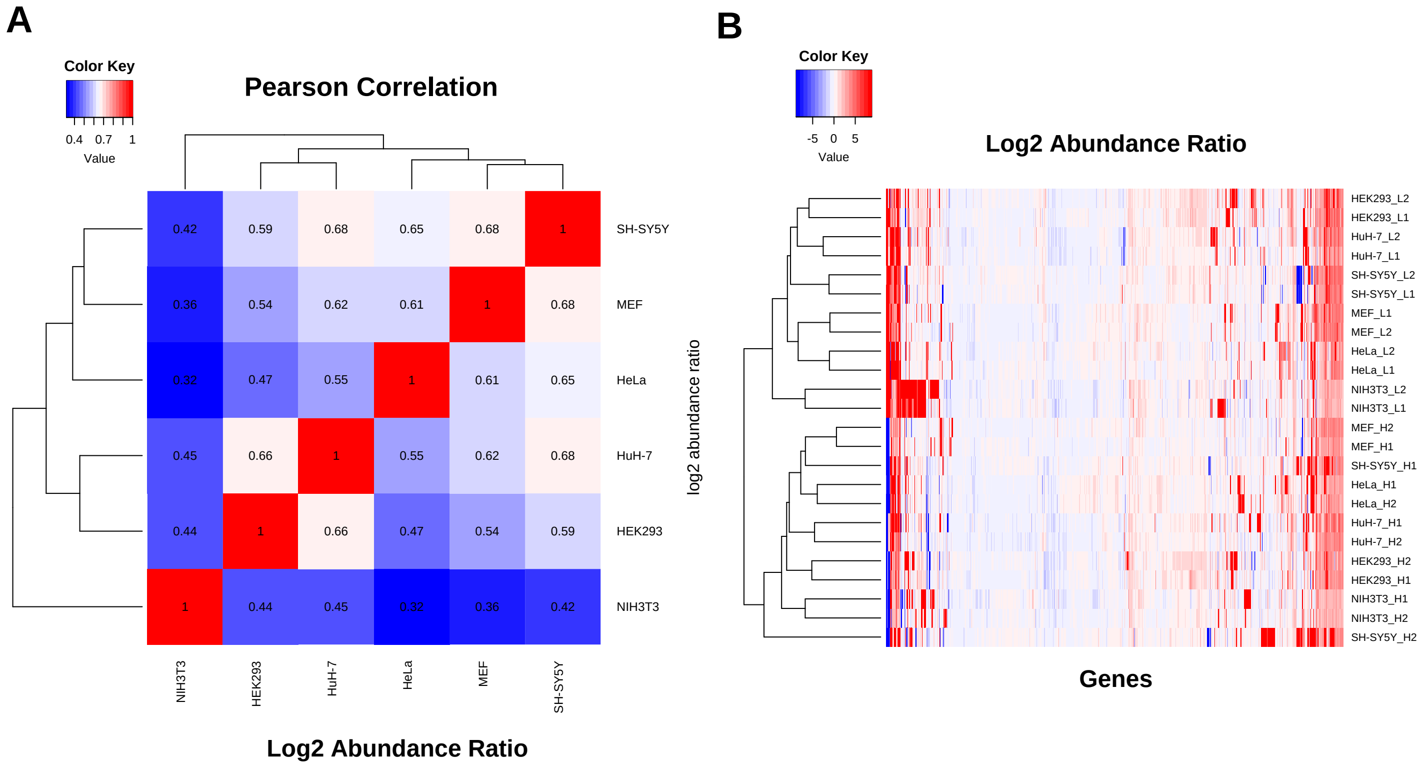


**Figure 2 - figure supplement 2: Analysis of cell type-specific protein expression profiles based on the datasets from lysosome-enriched fractions of the individual cell lines.** (**A**) Pearson correlation values of log2 abundance ratios (SPIONs/control) for binary comparisons of individual cell lines based on their mean values (n=4). (**B**) Heat map and hierarchical clustering of log2 abundance ratios (SPIONs/control). SPIONs: superparamagnetic iron oxide nanoparticles.

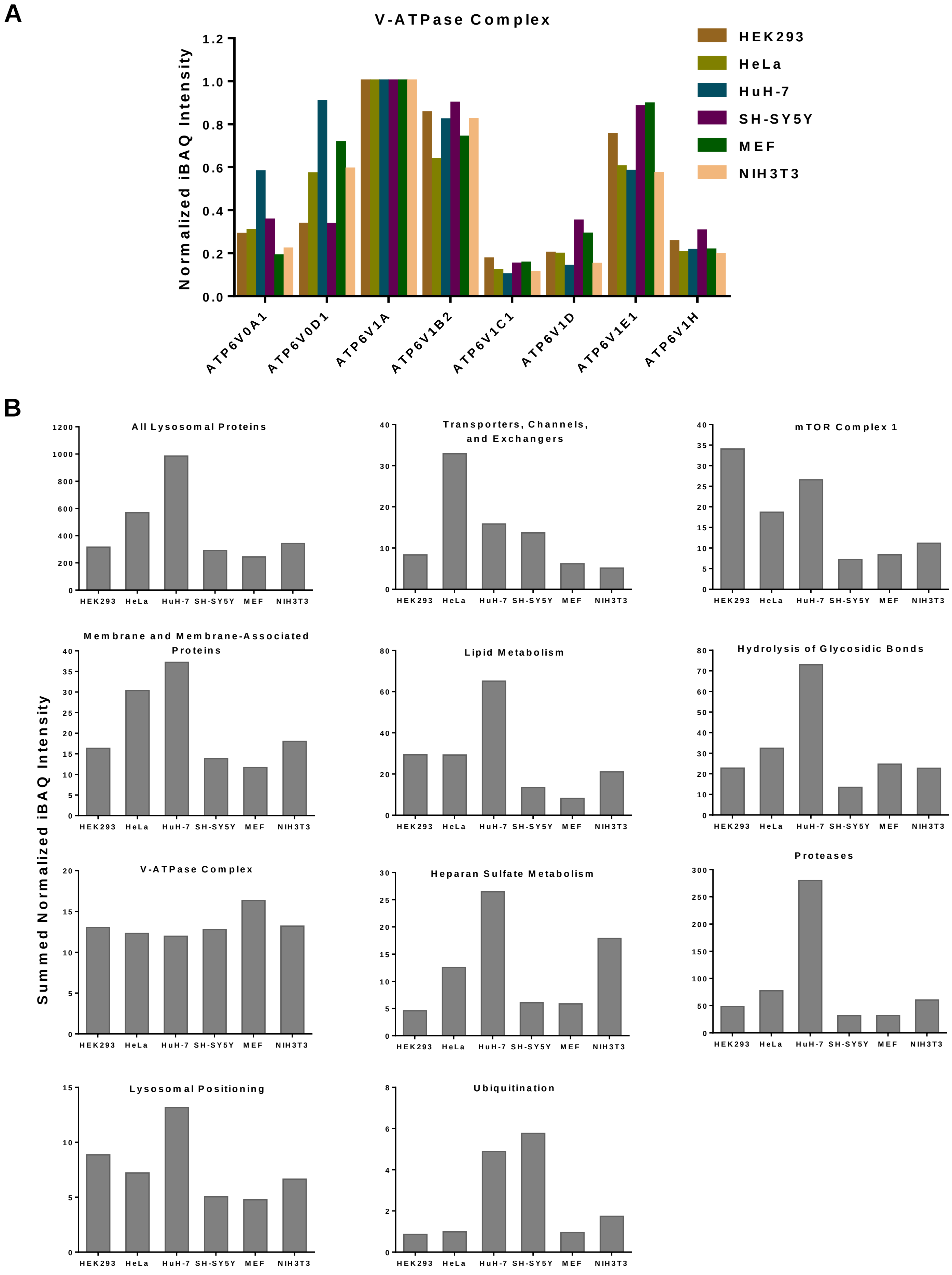
A
V-ATPase Complex
Normalized iBAQ Intensity
0.0
0.2
0.4
0.6
0.8
1.0
1.2
HEK293
HeLa
HuH-7
SH-SY5Y
MEF
NIH3T3
ATP6V0A1
ATP6V0D1
ATP6V1A
ATP6V1B2
ATP6V1C1
ATP6V1D
ATP6V1E1
ATP6V1H
B
Summed Normalized iBAQ Intensity
All Lysosomal Proteins
Transporters, Channels, and Exchangers
mTOR Complex 1
Membrane and Membrane-Associated Proteins
Lipid Metabolism
Hydrolysis of Glycosidic Bonds
V-ATPase Complex
Heparan Sulfate Metabolism
Proteases
Lysosomal Positioning
Ubiquitination
HEK293
HeLa
HuH-7
SH-SY5Y
MEF
NIH3T3

**Figure 4 – figure supplement 1: Analysis of protein abundance levels by intensity Based Absolute Quantification (iBAQ).** (**A**) Subunits of the V-ATPase complex which were detected in all cell lines with ≥ 10 unique peptides. Shown are the median iBAQ intensities (n = 4) normalized to the expression of ATP6V1A. (**B**) Summed normalized iBAQ intensities for distinct categories of known lysosomal proteins. iBAQ values of individual proteins were normalized to the median intensity of the eight V-ATPase subunits displayed in (A) in a replicate-wise manner. Subsequently, the mean of 4 biological replicates was calculated for each individual protein and the summed values were calculated. Only proteins present in ≥3 replicates of ≥2 cell lines were included.

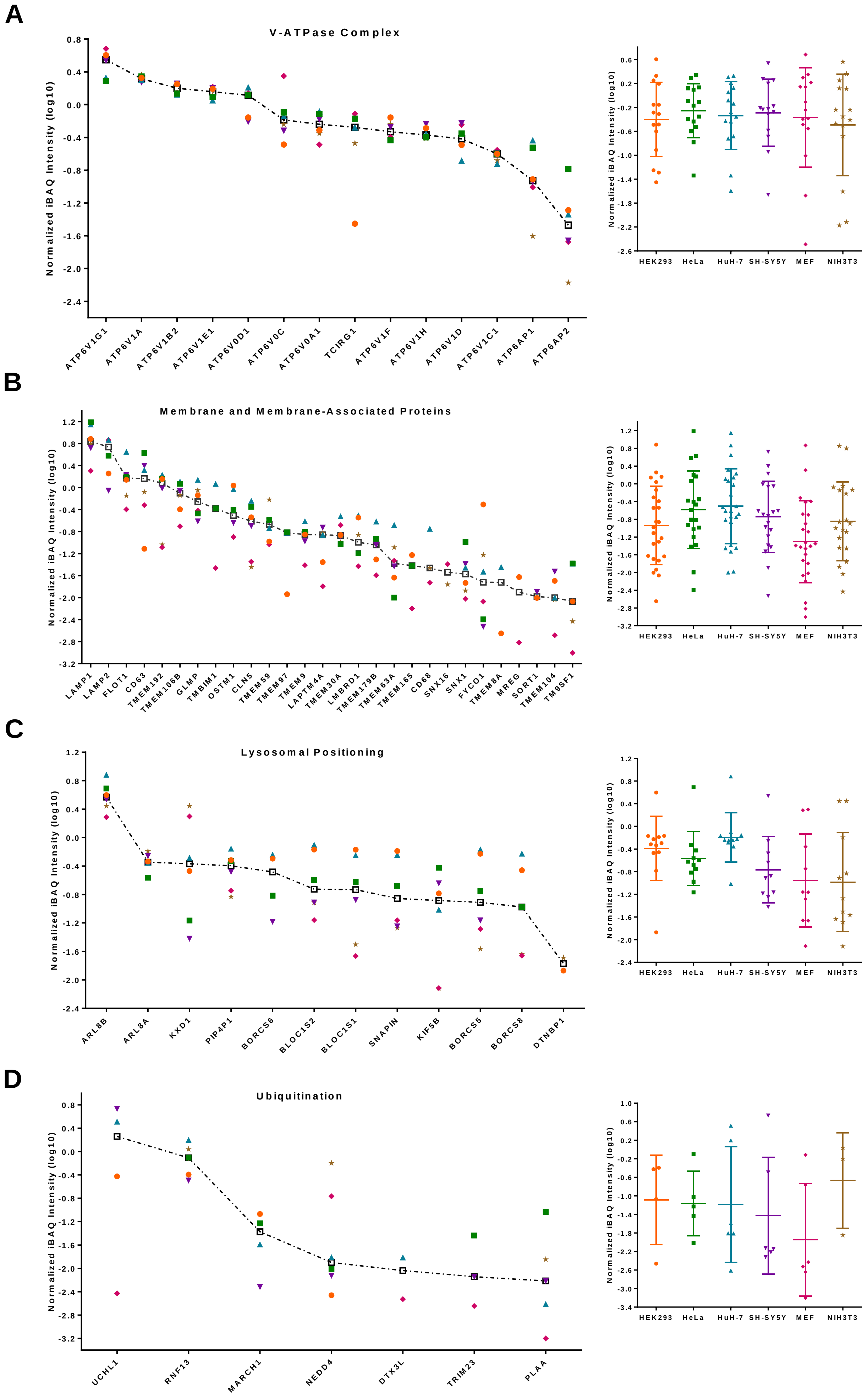
A
V-ATPase Complex
Normalized iBAQ Intensity (log10)
ATP6V1G1
ATP6V1A
ATP6V1B2
ATP6V1E1
ATP6V0D1
ATP6V0C
ATP6V0A1
TCIRG1
ATP6V1F
ATP6V1H
ATP6V1D
ATP6V1C1
ATP6AP1
ATP6AP2
HEK293
HeLa
HuH-7
SH-SY5Y
MEF
NIH3T3
B
Membrane and Membrane-Associated Proteins
Normalized iBAQ Intensity (log10)
LAMP1
LAMP2
FLOT1
CD63
TMEM192
TMEM106B
GLMP
TMBIM1
OSTM1
CLN5
TMEM59
TMEM97
TMEM9
LAPTM4A
TMEM30A
LMBRD1
TMEM179B
TMEM63A
TMEM165
CD68
SNX16
SNX1
FYCO1
TMEM8A
MREG
SORT1
TMEM104
TM9SF1
HEK293
HeLa
HuH-7
SH-SY5Y
MEF
NIH3T3
C
Lysosomal Positioning
Normalized iBAQ Intensity (log10)
ARL8B
ARL8A
KXD1
PIP4P1
BORCS6
BLOC1S2
BLOC1S1
SNAPIN
KIF5B
BORCS5
BORCS8
DTNBP1
HEK293
HeLa
HuH-7
SH-SY5Y
MEF
NIH3T3
D
Ubiquitination
Normalized iBAQ Intensity (log10)
UCHL1
RNF13
MARCH1
NEDD4
DTX3L
TRIM23
PLAA
HEK293
HeLa
HuH-7
SH-SY5Y
MEF
NIH3T3

**E**

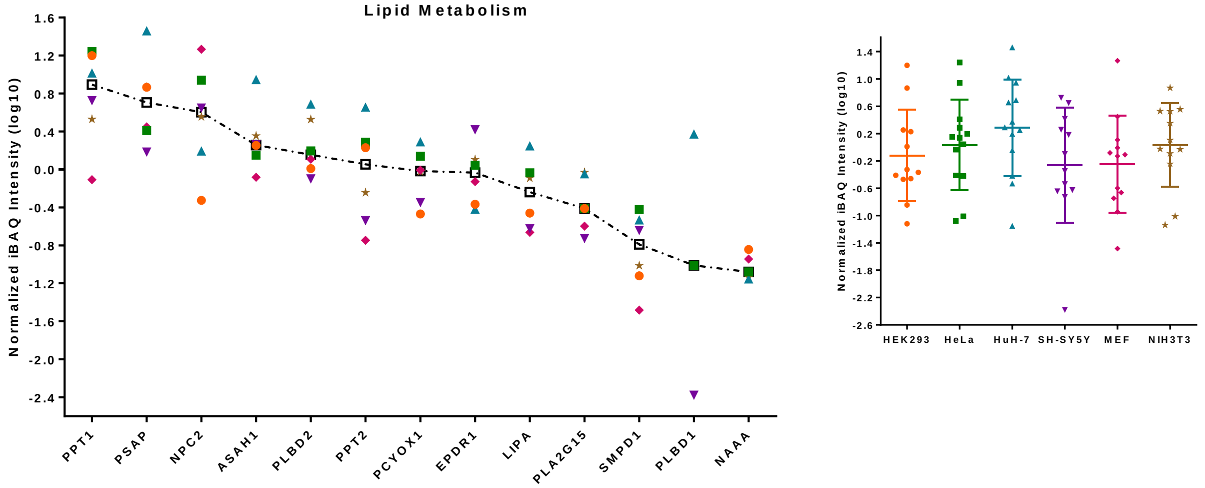


**F**

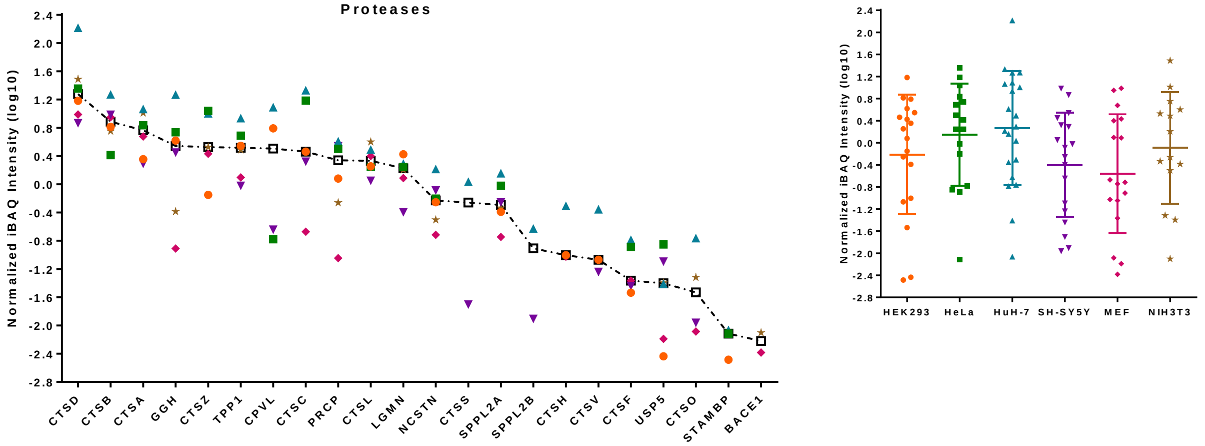


**G**

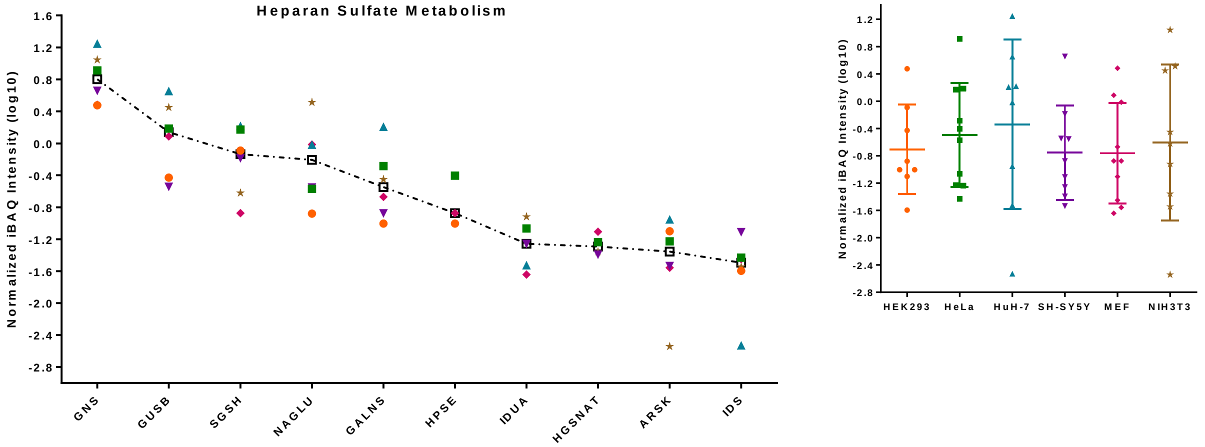


**Figure 4 - figure supplement 2: Individual V-ATPase normalized iBAQ values for proteins grouped in categories of similar function.** Shown are median V-ATPase normalized iBAQ values either for individual proteins sorted in descending order based on the median value of expression levels across all cell lines (scatter plots) or for the respective cell line (dotted box plots). (**A**) Proteins related to the V-ATPase complex. (**B**) Membrane and membrane-associated proteins. (**C**) Selected proteins known to play a role in lysosomal positioning, members of the BORC complex. (**D**) Lysosome-associated cytosolic ubiquitin ligases. (**E**) Proteins known to play a role in lysosomal lipid metabolism. (**F**) Lysosomal proteases. (**G**) Proteins involved in the degradation of heparan sulfate.

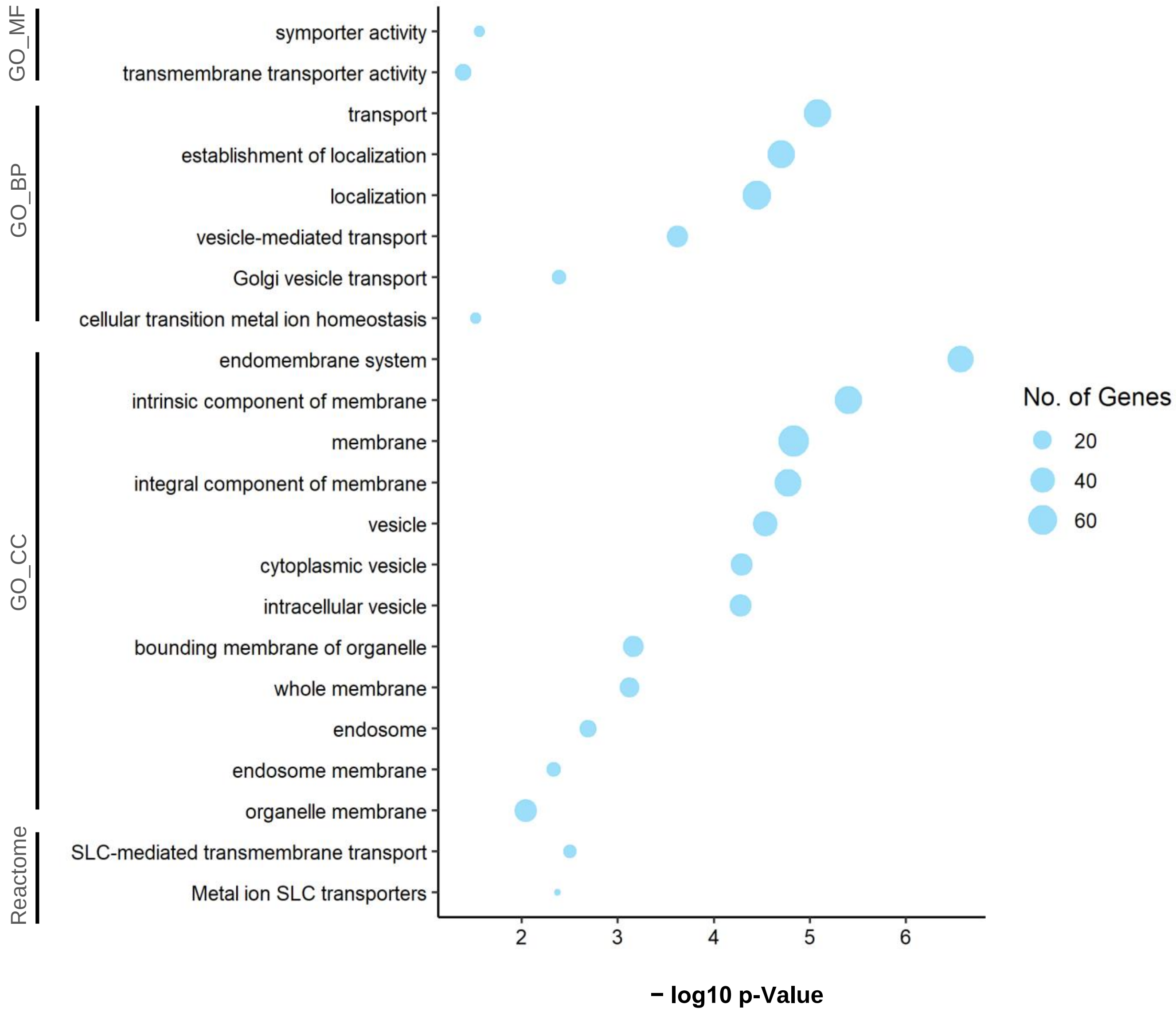


**Figure 6 – figure supplement 1: Gene Ontology analysis of high confidence potential novel lysosomal proteins.** Proteins which were identified to be significantly overrepresented in SPIONs receiving cells (p-value < 0.05) of ≥ 5 individual cell lines were subjected to GO and reactome pathway analyses. Bubble size correlates with the number of genes assigned to the respective category. GO: gene ontology; MF: molecular function; BP: biological process; CC: cellular component; SPIONs: superparamagnetic iron oxide nanoparticles.
